# A *Drosophila* model of oral peptide therapeutics for adult Intestinal Stem Cell tumors

**DOI:** 10.1101/2020.01.21.913806

**Authors:** Anjali Bajpai, Quazi Taushif Ahmad, Hong-Wen Tang, Nishat Manzar, Virender Singh, Ashwani Thakur, Bushra Ateeq, Norbert Perrimon, Pradip Sinha

**Affiliations:** Biological Sciences and Bioengineering, Indian Institute of Technology Kanpur, Kanpur, India; Department of Genetics, Harvard Medical School, Boston, MA 02115, USA; Howard Hughes Medical Institute, Boston, MA 02115, USA; Department of Physiology & Biophysics, Case Western Reserve University, Cleveland, OH 44106

**Keywords:** Drosophila, Peptide therapeutic, Yki, Intestinal Stem Cells, Integrin signaling

## Abstract

The proto-oncogene YAP /Yki, a transcription co-factor of the Hippo pathway, has been linked to many cancers. YAP interacts with DNA-binding TEAD/Sd proteins to regulate expression of its transcriptional targets. Disruption of YAP-TEAD therefore offers a potential therapeutic strategy. The mammalian Vestigial Like (VGLL) protein, specifically its TONDU domain, has been shown to competitively inhibit YAP-TEAD interaction and a TONDU peptide can suppress YAP-induced cancer. As TONDU could potentially be developed into a therapeutic peptide for multiple cancers, we evaluated its efficacy in Yki-driven adult Intestinal Stem Cell (ISC) tumors in Drosophila. We show that oral uptake of the TONDU peptide is highly effective at inhibiting Yki-driven gut tumors by suppressing YAP-TEAD interaction. Comparative proteomics of early and late stage Yki-driven ISC tumors revealed enrichment of a number of proteins, including members of the integrin signaling pathway, such as Talin, Vinculin and Paxillin. These, in turn displayed a decrease in their levels in TONDU-peptide treated tumors. Further, we show that Sd binds to the regulatory region of integrin-coding gene, mew, which codes for αPS1, a key integrin of the ISCs. In support to a possible role of integrins in Yki-driven ISC tumors, we show that genetic downregulation of *mew* arrests Yki-driven ISC proliferation, reminiscent of the effects of TONDU peptide. Altogether, our findings present a novel platform for screening therapeutic peptides and provide insights into tumor suppression mechanisms.

**SIGNIFICANCE STATEMENT:** Discovering novel strategies to inhibit oncogene activity is a priority in cancer biology. As signaling pathways are widely conserved between mammals and Drosophila, these questions can be effectively addressed in this model organism. Here, we show that progression of *Drosophila* Intestinal Stem Cell (ISC) tumors induced by gain of an oncogenic form of the transcription co-factor Yki can be suppressed by feeding a peptide corresponding to the conserved TONDU domain of Vestigial (Vg), which blocks binding of Yki to the Sd transcription factor. Further, we show that down regulation of the integrin signaling pathway is causally linked to TONDU-peptide-mediated ISC tumor suppression. Our findings reveal that *Drosophila* can be successfully used to screen peptides for their therapeutic applications.

## INTRODUCTION

Drosophila has emerged as an effective cancer model for the screening of small molecule therapeutics (1–4). Of interest, *Drosophila* adult gut tumors, such as Yki-driven intestinal stem cell (ISC) tumors (5) or a multigenic hindgut model of colon cancer (6), have been successfully used to screen for anti-proliferative small molecules. While cancer-promoting rogue kinases are amenable to inhibition by small molecules, others, such as transcription factors and co-factors, are largely considered undruggable (7, 8). In this regard, peptides are particularly attractive as therapeutic molecules (9–11) because of their high selectivity, improved tolerance and ability to target large interacting interfaces (12).While most peptide therapeutics require parenteral injection (10), their oral delivery is highly desirable and currently a number of orally derived therapeutic peptides are being tested in clinical trials (10).

The proto-oncogene YAP (Yes-associated protein), a transcription co-factor of the Hippo pathway, has been linked to many cancers (see review (13)). YAP interacts with DNA-binding TEAD proteins (transcriptional enhanced associate domain, TEAD1-4) to regulate expression of its transcriptional targets, and an increase in levels of TEAD proteins has been observed in a wide range of human cancers (14). YAP binds to TEAD via an unusually large interface, the Ω-loop (12, 15) that lacks a defined binding pocket, making it an unlikely target of inhibition by small molecules. TEAD proteins also interact with other transcriptional co-factors, such as the Vestigial Like (VGLL1-4) proteins, via the conserved 26 amino acid TONDU domain present in VGLL proteins (15). Further, VGLL proteins behave as tumor suppressors in mammals due to their ability to inhibit YAP-TEAD interactions, as seen in lung (16) and breast (17) carcinomas. Interestingly, a synthetic peptide analog of the TONDU domain of VGLL4 was found to inhibit gastric cancer growth (18).

Yorkie (Yki), the *Drosophila* homolog of mammalian YAP, was first identified as a nuclear effector that triggers epithelial proliferation upon deregulation of Hippo signaling (19). Subsequent studies revealed its role as a developmental regulator of organ growth (20) and as an oncogene (21, 22). Like its mammalian counterpart, *Drosophila* Yki binds to a TEAD domain-containing protein, Scalloped (Sd) (23, 24), which can also bind to Vestigial (Vg) (25, 26), the *Drosophila* counterpart of mammalian VGLLs. *Drosophila* Vg protein was shown to possess the TONDU domain that mediates its interaction with Sd (27).

The similarities between mammalian VGLLs-TEADs-YAP and *Drosophila* Vg-Sd-Yki suggest that a TONDU peptide could suppress Yki-driven *Drosophila* tumors by competitively inhibiting the Yki-Sd interaction. Further, use of the well characterized Yki-driven ISC tumor model as a platform for peptide therapeutics also holds the promise to unravel genetic network that drives ISC tumor progression and, conversely, can be suppressed to restrain tumor growth. Further, Here, we used the adult *Drosophila* gut, where Sd-dependent Yki activity is required for ISC homeostasis (28–31), to test whether the TONDU peptide can suppress unrestricted ISC proliferation associated with expression of an activated form of Yki (32, 33). We show that adult flies displaying ISC-specific gain of activated Yki fail to display robust ISC tumors when they are raised in food supplemented with the TONDU peptide. Comparative proteome analysis of Yki-driven ISC tumors and those from flies fed with TONDU, revealed perturbations in integrin-associated proteins, suggesting that they could play a critical role in Yki-driven ISC tumors. In support of this hypothesis, downregulation of integrin αPS1 inhibits Yki-driven ISC tumorigenesis. These findings reveal that *Drosophila* ISC tumor models can be used to screen for anticancer peptides and to unravel mechanisms of tumor suppression.

## RESULTS AND DISCUSSION

### Genetic suppression of Yki-driven ISC tumor growth by the TONDU peptide

The *Drosophila* adult gut is made up of three cell types: differentiated enterocytes (ECs), entero-endocrine cells (EEs), and intestinal stem cells (ISCs) (Figure 1A-B, (34)). Expression of a phosphorylation-defective and therefore constitutively active form of Yki in the ISCs (*esg-Gal4 Gal80^ts^>*UAS-yki*^3SA^*, referred to as *esg^ts^>yki^3SA^*) triggers ISC over-proliferation (32) (Figure 1C, Figure S1A), as revealed by increased 5-ethynyl-2 deoxyuridine (EdU) uptake (Figure S1B), Phospho-Histone H3 (PH3) staining (Figure S1C), and elevated expression of matrix metalloproteinase genes (MMPs) (Figure S1D). Further, as previously reported (32), aged *esg^ts^>yki^3SA^*flies display tumor-associated systemic wasting syndrome, which is characterized by abdominal bloating (Figure 1H), organ atrophy (Figure S1E), and elevated levels of the insulin antagonist ImpL2 (Figure 1J) (32, 35). We also observed an increase in the transcript levels of Yki targets such as *myc, cycE, diap1and exp* (Figure 1J) in *esg^ts^>yki^3SA^* tumors. Further, we note that *esg^ts^>yki^3SA^* tumors show concomitant increase in Sd (Figure 1D, J), the DNA-binding partner of Yki.

**Figure 1.**
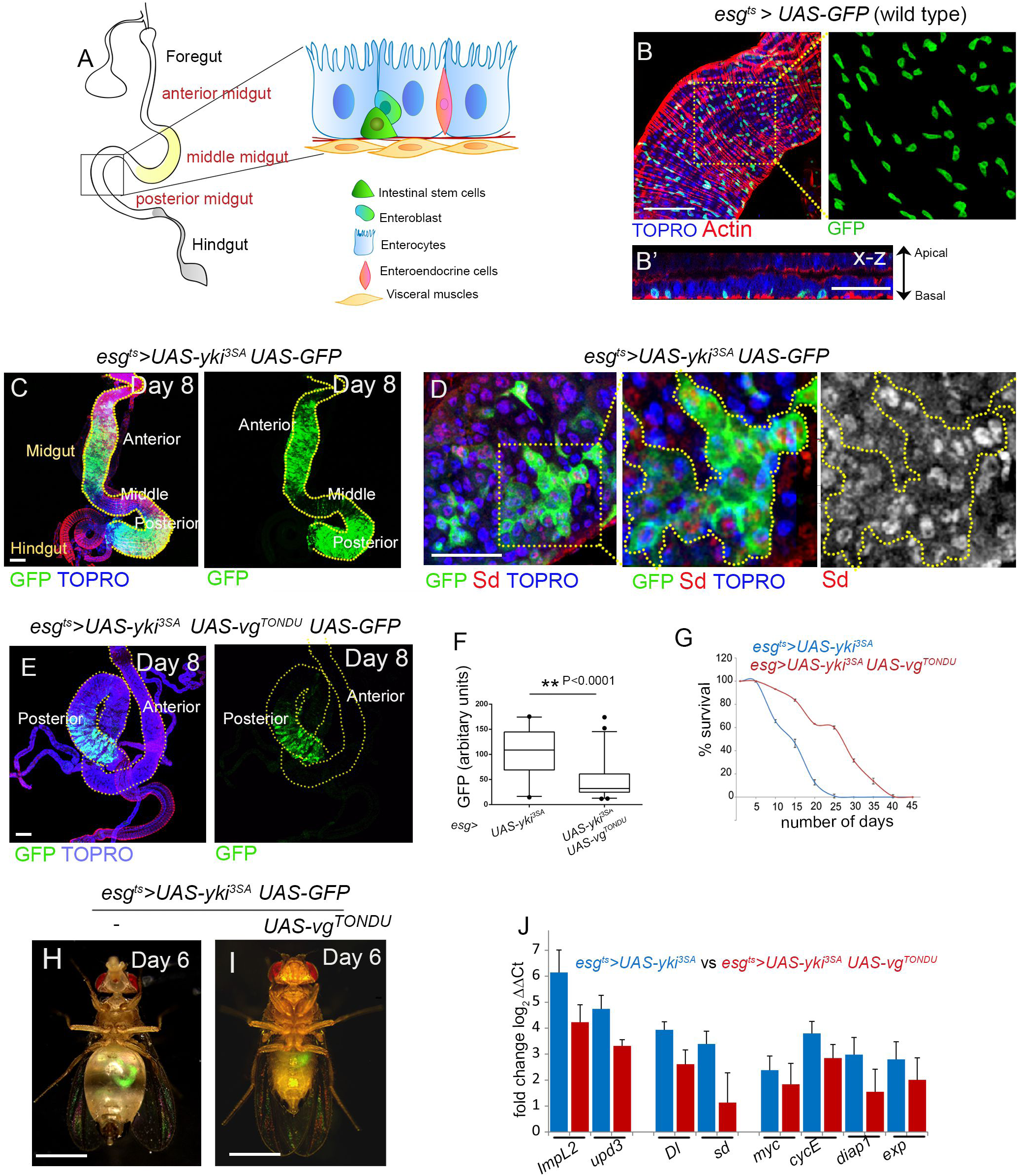
Expression of the TONDU peptide inhibits Yki-driven ISC tumors. (A) Schematic representation depicting the different cell types in the adult *Drosophila* gut. (B, B’) *esg^ts^>UAS-GFP* labels ISCs in the *Drosophila* midgut. (B) ISCs (marked by GFP) are interspersed throughout the gut. Overlying muscles are marked with F-Actin (red). (B’) x-z section displaying basally located ISCs (GFP). (C) *esg^ts^>yki^3SA^*UAS-GFP gut shows an increase in ISC numbers. (D) *esg^ts^>yki^3SA^*UAS-GFP tumors show increase in Sd level. (E) Decrease in ISCs (marked by GFP) in the anterior and posterior midgut of *esg^ts^>yki^3SA^ UAS-vg^TONDU^* flies that coexpress TONDU peptide. (F) Quantification of GFP in TONDU-expressing and non-expressing *esg^ts^>yki^3SA^*guts. (G) Increase in survival of *esg^ts^>yki^3SA^*UAS-vg^TONDU^ UAS-GFP flies (n=50) compared to *esg^ts^>yki^3SA^* UAS-GFP flies. (H) Abdominal bloating in *esg^ts^>yki^3SA^*UAS-GFP flies as seen on day 6 after tumor induction (n=19/25 are bloated). (I) *esg^ts^>yki^3SA^ UAS-vg^TONDU^UAS-GFP* flies display delay in bloating (n=14/25 are not bloated). (J) qPCR displaying the decrease in mRNA levels of candidate genes in TONDU-expressing flies. Data presented as mean ±SE. Scale bars 100 µm in all, except B’ and D: 50 µm, and H and I: 1mm.

To test whether the fly equivalent of the 25 amino acid long TONDU domain of Vg (CVVFTNYSGDTASQVDEHFSRALNY) can arrest neoplastic tumor progression similar to the full length protein observed earlier (21), we examined somatic clones in wing imaginal discs that lack the tumor suppressor *lethal giant larvae (lgl)* and that express an activated form of Yki (*UAS-yki^S168A^* (36), referred to as lgl *UAS-yki*) and the TONDU peptide. Strikingly, marked reduction in the growth of lgl *UAS-yki UAS-vg^TONDU^* tumors was observed as compared to their lgl *UAS-yki* counterparts (Figure S2A, B).

Next, we examined the effect of TONDU expression on Yki-induced ISC tumors in the adult midgut. Co-expression of TONDU and activated Yki (*esg^ts^>yki^3SA^ UAS-vg^TONDU^*) revealed that TONDU could inhibit Yki-driven ISC proliferation throughout the midgut, an effect that was more striking in the anterior midgut (Figure 1E). Consistent with these observations, we also noted a marked reduction in the number of GFP-marked ISCs (Figure 1F), with an accompanying decrease in their proliferation, marked by EdU uptake (Figure S2C-E). In addition, TONDU-expressing *esg^ts^>yki^3SA^* flies displayed improved life span (Figure 1G) and delayed onset of tumor-associated organ wasting phenotypes (Figures 1I and S1F). Consistent with reduced abdominal bloating, TONDU-expressing tumor bearing flies displayed a decrease in hemolymph content (Figure S2F), improved muscle activity (Figure S2G) and decreased levels of Impl2 (Figure 1J) in addition to other transcriptional targets of Yki (Figure 1J). By contrast, gain of the TONDU peptide alone in ISCs (*esg^ts^>vg^TONDU^*) failed to alter the number of ISCs (Figure S2H, I). Altogether, these results reveal that Yki-driven ISC tumors are suppressed upon co-expression of the TONDU peptide, with an accompanying delay in the onset of tumor-associated syndromes.

### Oral uptake of synthetic TONDU peptide inhibits Yki-driven ISC tumor

Next, we tested whether a synthetic TONDU peptide could inhibit Yki-driven ISC tumors akin to endogenously expressed peptide. We fed adult flies with varying concentrations of TONDU peptide linked to an HIV-TAT motif (RKKRRQRRR) and a nuclear localizing signal (NLS) (Figure 2A) to facilitate its cellular uptake (37) and nuclear localization, respectively. This TAT-NLS-TONDU peptide is referred to as TONDU peptide in subsequent part of the text. Prior to its oral administration to adult flies we first confirmed cellular uptake of the fluorescent-labeled TONDU peptide in S2R+ cells, as observed by its cytoplasmic and nuclear localization (Figure 2B). Further, to test whether the TONDU peptide can inhibit Yki-Sd complex formation, we used the Hippo-response-element (HRE)-luciferase reporter (23), which serves as a readout for Yki-Sd transcriptional activity. Specifically, we co-transfected S2R+ cells with the reporter along with Yki and Sd-expressing vectors, then treated the cells with 100 nM of TONDU peptide, and observed a moderate but consistent decrease in luciferase activity (Figure 2C). Next, to confirm binding of the synthetic TONDU peptide to Sd and subsequent inhibition of Yki-Sd interaction, we carried out co-immunoprecipitation studies using FLAG-tagged TONDU peptide in S2R^+^ cells transfected with HA-Sd and GFP-Yki. Co-immunoprecipitation experiments revealed that the Yki-Sd interaction is indeed significantly reduced upon incubating S2R+ cells with TONDU for 24 hours (Figure 2D). Finally, when purified HA-Sd from S2R+ cells was incubated with FLAG-tagged TONDU peptide, TONDU displayed binding with Sd, as revealed by immunoblots using anti-Flag antibody (Figure 2E). Together, these results demonstrate that TONDU can disrupt the Sd-Yki interaction by binding to Sd.

**Figure 2.**
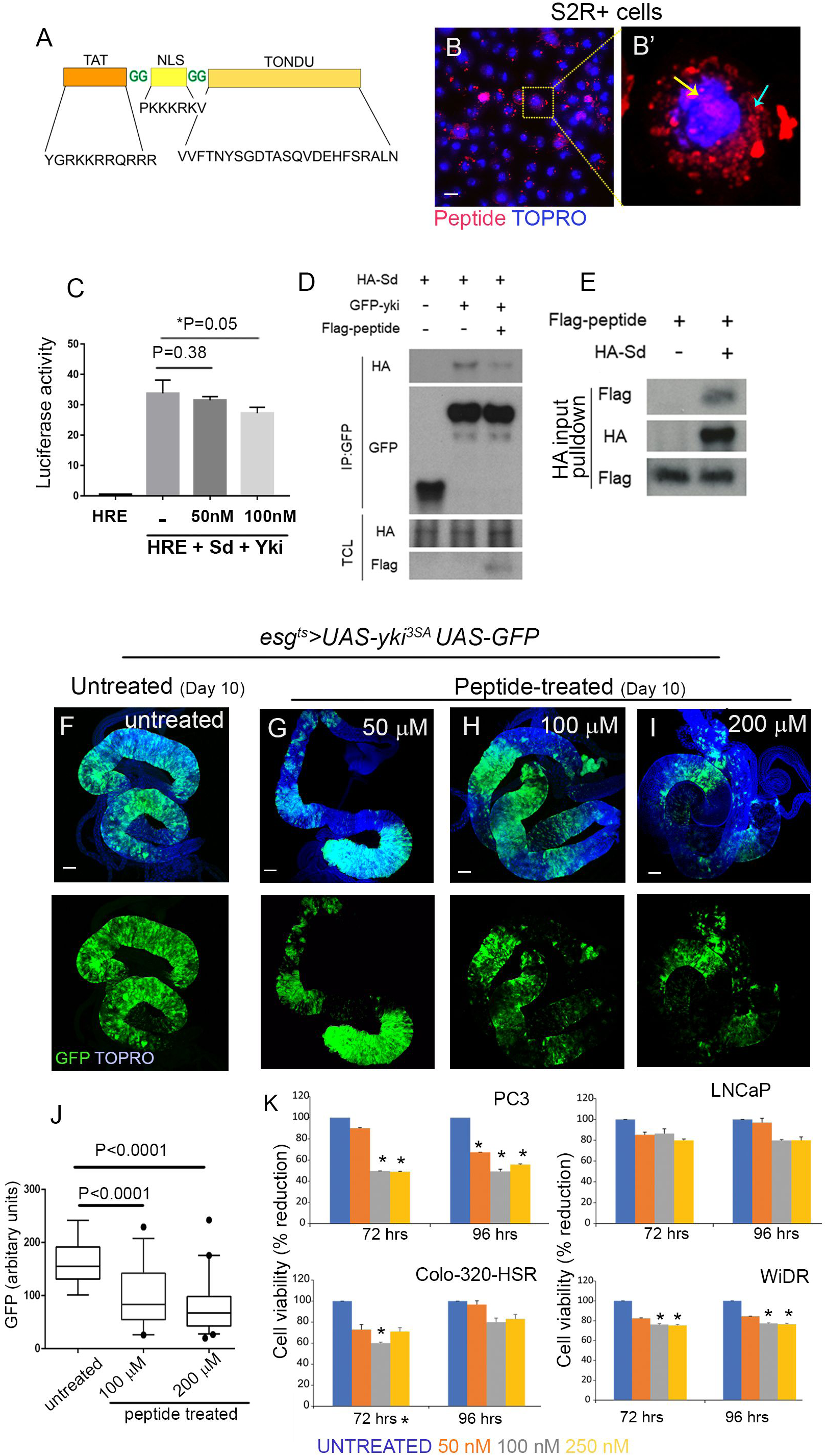
Synthetic TONDU peptide inhibits Yki-driven ISC tumors. (A) Representation of the synthetic TONDU peptide. (B-B’) Nuclear localization of fluorescent-tagged (red) TONDU peptide in S2R+ cells. (B’) Magnified view of the boxed area in B. TONDU-Peptide (red) in the nucleus (yellow arrow) and cytoplasm (blue arrow). (C) Decrease in HRE-Luciferase reporter activity in S2R+ cells when treated with TONDU peptide. (D-E) Immuno-blots showing competitive binding of TONDU peptide to Yki-Sd complex (D). (E)Binding of TONDU peptide to Sd. (F-I) Guts from *esg^ts^>yki^3SA^* flies fed on TONDU peptide. (F) Unfed (control), (G) 50 μM (n=10), (H) 100 μM (n=12) and (I) 200 μM (n=10). (J) Quantification of GFP in TONDU peptide-fed and -unfed *esg^ts^>yki^3SA^* flies. (K) Viability of cancer cells on treatment with TONDU peptide, as estimated using the Resazurin cell viability assay. Scale bars: 10µm in B; 100 µm in F-I.

We next tested whether oral uptake of TONDU peptide could inhibit *esg^ts^>yki^3SA^* tumors. To do this, *esg^ts^>yki^3SA^* flies were collected 24 hours post eclosion and fed for ten days on food supplemented with TONDU peptide at a final concentration of 50, 100 or 200 µM. We noted a progressive reduction in tumor mass (Figure 2F-I), marked by a decrease in the number of GFP-marked ISCs (Figure 2J) with increasing concentration of TONDU peptide in the food. By contrast, the tumor load was only moderately reduced when these flies were fed with a sequence-scrambled TONDU peptide (Figure S3A) at comparable concentrations (see Figure S3B-F); this residual inhibition of ISC proliferation is presumably due to a partial retention of the secondary structures necessary for TONDU activity (15) in the scrambled-TONDU peptide (Figure S3G). Further, to confirm cellular uptake of TONDU peptide by the gut epithelia, we fed FLAG-tagged TONDU peptide (at final concentration of 200 µM) to *esg^ts^>yki^3SA^* flies, and detected its cellular uptake in gut lysates followed by immunoblotting using the anti-FLAG antibody (Figure S3H). In parallel, we also noted that feeding TONDU (at 200 µM) did not affect the numbers of ISCs in control (*esg^ts^*>GFP) guts (Figure S3I). In addition, two cell lines derived from human tumors with elevated *YAP1* levels (Figure S3J), PC3 (prostate cancer cells) and COLO-320 (colorectal cancer cells), displayed growth arrest upon uptake of TONDU peptide (Figure 2K), whereas a cell line with negligible levels of *YAP1* (Figure S3J), LNCaP, did not (Figure 2K). Altogether, these results suggest that TONDU is therapeutically relevant in a number of YAP-driven tumors irrespective of their tissues of origin.

### Yki-driven tumor proteome reveals enrichment in integrin pathway

We reasoned that proteins that are significantly perturbed in *esg^ts^>yki^3SA^* tumors and restored to normal levels following TONDU feeding may represent critical Yki-Sd targets that are critical to ISC tumorigenesis, and, therefore, could be therapeutically relevant. Thus, we carried out a proteome analysis using unlabeled LC-MS/MS of *esg^ts^>yki^3SA^* tumors on day 1 and day 7 of tumor induction, with or without TONDU peptide supplementation of the food. Altogether, we identified 1219 proteins (including isoforms), corresponding to 2771 unique Uniprot IDs at an FDR cutoff of q<0.05 (Figure 3A, Table S1). We next compared the proteomes of *esg^ts^>yki^3SA^* tumors on day 7 versus day 1 of tumor induction, and prioritized those proteins which displayed at least log_2_ ±2 fold change (at a p value <0.05) for further analysis. Fold change was derived from the abundance measure of peptides (for a given protein) in day 7 versus day 1 of *esg^ts^>yki^3SA^* tumors (See SI methods) and a list was generated of 127 differentially detected proteins (corresponding to 144 unique Uniprot IDs, including isoforms) that matched to 55 unique genes (Figure 3B and Table S2). For 45 of the 55 genes, the gene products showed a >log_2_ 2 fold increase while 10 displayed <log_2_ 2 fold decrease in their protein levels in day 7 *esg^ts^>yki^3SA^* tumors (Figure 3B and Table S2).

**Figure 3.**
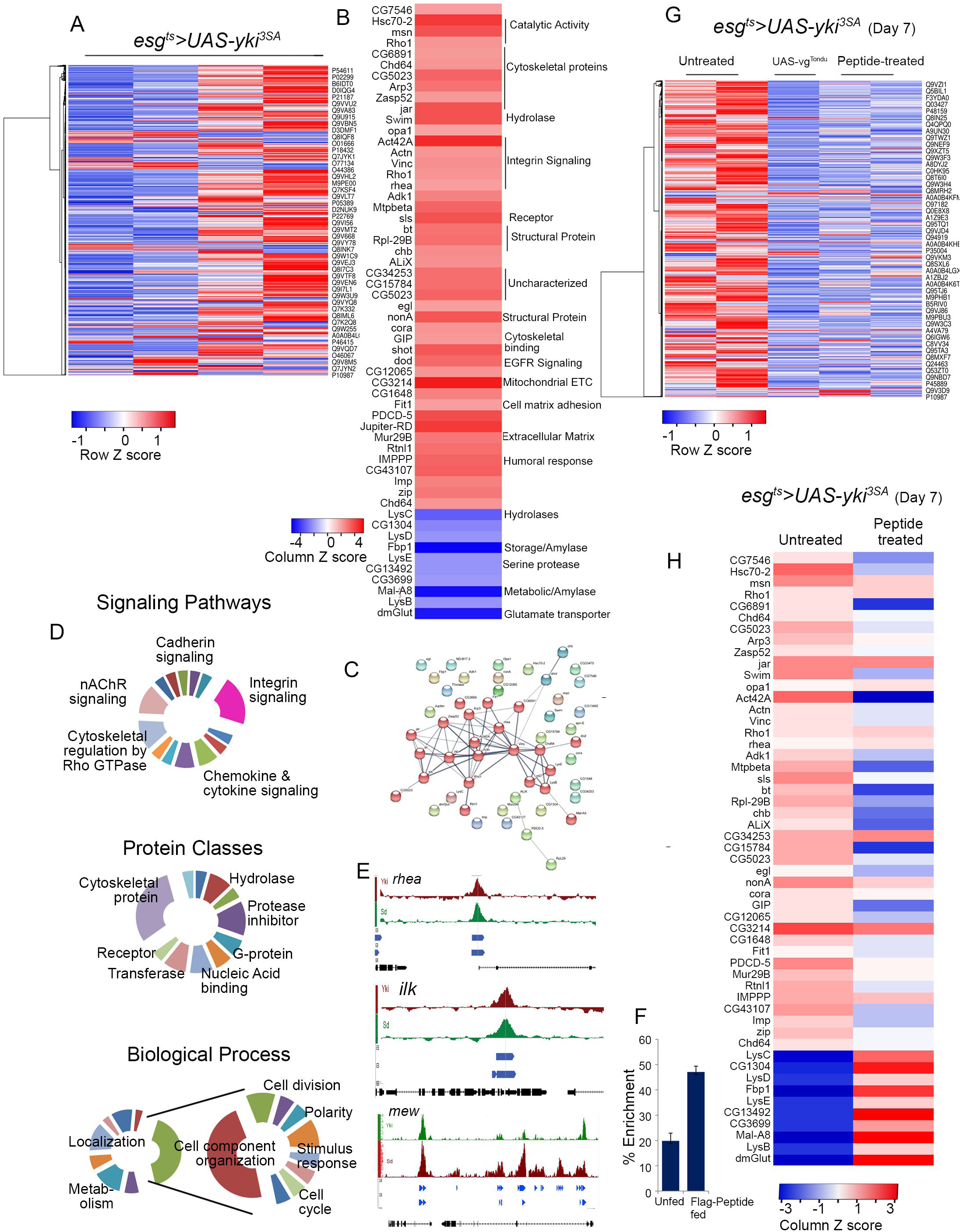
Comparative proteomic analysis of Yki-driven ISC tumors and tumors inhibited by the TONDU peptide: (A) Heat map displaying changes in protein levels in day 7 and day 1 of *esg^ts^>yki^3SA^* tumors. (B) 55 differentially (>±2 log_2_fold, *p*=0.05) expressed proteins in day 7 *esg^ts^>yki^3SA^* tumors. (C) Protein-protein interaction (PPI) network of enriched proteins (>log_2_ 2 fold) in *esg^ts^>yki^3SA^* tumors generated with STRING (38) representing 55 nodes and 63 edges (PPI enrichment *p*<0.0001). (D) Different Gene Ontology classes identified by PANTHER (58) in differentially expressed proteins between *esg^ts^>yki^3SA^*-day7 versus -day 1 tumor proteome. (E) Sd and Yki binding sites in the regulatory regions of select integrin pathway members as determined in (44). (F) Percent enrichment for Sd-binding upstream of *mew* (αPS1) inferred by ChIP with anti-FLAG antibody. (G) Heatmap displaying the effect of TONDU peptide on *esg^ts^>yki^3SA^* tumor proteome. (H) Heat map displaying change in levels of protein (>±2 fold in day 7 tumors) upon TONDU peptide treatment.

To determine whether these enriched proteins in the ISC tumors have a significant biological association we performed a protein-protein interaction (PPI)-network analysis using STRING (38), which revealed significant (*p*<0.001) interaction among some of the enriched proteins (Figure 3C). We note that the enriched gene-set include the secreted Wingless (Wg) transporter Swim (39) and known members of the Hippo protein-protein interaction network (40), including the junction proteins Coracle, Jar, and Misshapen (Table S3). Furthermore, comparison of the *esg^ts^>yki^3SA^* day 7 proteome with the recently published transcriptome (33) of *esg^ts^>yki^3SA^* tumors of comparable age revealed a close correlation between changes in proteins and their respective transcript levels (r=0.548) (Figure S4A).

Next, to look for signaling pathways that were perturbed in Yki-driven ISC tumors, we undertook a gene ontology (GO) classification of the genes enriched in *esg^ts^>yki^3SA^* tumors. GO classification revealed perturbations in several signaling pathways and protein classes (Figure 3D, Table S4). In particular, we observed an increase in protein levels of key members of the integrin signaling pathway, including Talin (2.39 log_2_fold), Talin-interacting adaptor proteins, Vinculin (2.4 fold), and Paxillin (6.05 fold). Other members, such as αPS3 and integrin-linked kinases, also displayed about a 2-fold change, albeit at *p*>0.05 (Table S5). Consistent with these findings, a concomitant increase in the transcript levels of these proteins was observed in the *esg^ts^>yki^3SA^* transcriptome (33) (Table S5). Further, many integrin pathway components, including integrins αPS1, αPS2, αPS3 and βPS as well as integrin-binding ligands LamA and LamB (33), which were not detected in our proteomic study, were also found to be transcriptionally upregulated in the RNA-Seq data (Table S5). Proteome comparison between day 7 and day 1 *esg^ts^>yki^3SA^* tumors also displayed significant increase in protein levels of polarity proteins such as tight junction protein, Ferritin, Fit1, and the apical protein Shot (Figure 3B), which are known to be regulated by integrin signaling (41). We note that enrichment of integrin pathway members and associated proteins, as revealed in the tumor proteome, could be of functional relevance in Yki-driven ISC tumorigenesis, since many integrin pathway members are reported to be enriched in ISCs (42), and play an essential role in ISC survival (42) and maintenance of epithelial polarity in the gut (41).

To further determine whether the genes enriched in *esg^ts^>yki^3SA^* tumors could be Yki-Sd transcriptional targets, we searched for putative Yki-Sd binding sites in their upstream regulatory region. Chromatin binding studies have previously revealed genome-wide binding of Yki (43, 44) and Sd (44) upstream of many of their transcriptional targets. We noted that ∼51% (23 of 45) of the genes enriched in *esg^ts^>yki^3SA^* tumors displayed putative Sd and Yki binding sites in their upstream regulatory regions based on earlier binding studies (44) (Table S6). Interestingly, several key members of the integrin pathway, including mew, which codes for integrin αPS1, the adaptor proteins *vinculin* and *paxilin*, and *integrin-linked kinase*, displayed Sd binding (44) (Figure 3E, Table S6), suggesting their possible transcriptional regulation by the Yki-Sd complex. We therefore examined the binding of Sd to the upstream regulatory region of mew, as its protein product is the most abundant integrin in the ISCs (42), and performed chromatin immuno-precipitation using anti-FLAG antibody on gut lysate of *esg^ts^>yki^3SA^* flies fed on FLAG-tagged TONDU peptide. qPCR was done to determine the abundance of the putative genomic binding sites in the pull-down fraction. We observed 47.04% (SD=2.3) enrichment of a putative Yki-Sd binding site in the FLAG-enabled pull-down fraction in the TONDU peptide fed flies (Figure 3F), compared to 19.8% (SD=3.0) enrichment in gut lysate from flies raised on control food, suggesting that *mew* transcription is most likely regulated by Sd in the ISCs.

We next compared the gut proteomes of *esg^ts^>yki^3SA^* flies fed on TONDU peptide-supplemented food and those displaying co-expression of the TONDU peptide (*esg^ts^>yki^3SA^*UAS-vg ^TONDU^), to *esg^ts^>yki^3SA^* flies of comparable age (day 7) raised on normal food (Figure 2G, H). In particular, we examined the status of proteins in TONDU-peptide-treated proteome that were ±>log_2_ 2 fold perturbed in untreated tumors (Figure 3B and Table S2). We observed an overall decrease in the protein levels of perturbed genes in peptide-fed tumors (Figure 3G), which coincided with a decrease in protein levels of genes involved in generic cellular processes such as RNA processing. Proteins encoded by genes such as *Pre-RNA processing factor 19 (Prp19)* (−2.16 fold) (45) and *rumpelstiltskin (rump)* (−3.15 fold) were notably downregulated. Further, proteins enriched in the tumors at day 7 were mostly down-regulated upon peptide treatment (Figure 2H, Table S7). These included many members of the integrin pathway, such as Paxillin (−1.9 fold), Vinculin (−1.3 fold) and Talin (−1.2 fold). In addition, we also observed a decrease (−2.3 fold) in mitochondrial trifunctional protein β (Mtp-β), which catalyzes oxidation of long chain fatty acids (46), a possible energy source for tumors (47). We also note a decrease in peptide-treated tumors in proteins such as Chromosome bows (Chb) (−2.16 fold) that are involved in mitotic spindle assembly (48), an effect that could contribute to the observed decrease in cell proliferation of the ISC (Figure 2G-I). Altogether, these analyses reveal that TONDU peptide-treated ISC tumors display down regulation of integrin signaling components additional to those that are recruited for cellular processes such as RNA regulation (45), energy homeostasis (46) and mitosis (48) (Table S7).

### Genetic suppression of integrin signaling phenocopies TONDU-mediated suppression of Yki-driven ISC tumors

Integrins form an essential component of the *Drosophila* gut epithelia, including the basally located ISCs (41, 42) (Figure 4A, B). Consistent with the enrichment of integrin pathway members in *esg^ts^>yki^3SA^* proteome, we observed an overall increase in membrane localization of integrin αPS1 (Figure 4A) and Talin (Figure 4B) in *esg^ts^>yki^3SA^* tumors. This observation, together with our findings that Yki-Sd bind upstream of *mew* and that suppression of integrin pathway members in peptide fed tumors, suggests that integrin down-regulation may mediate the effect of TONDU peptide inhibition of *esg^ts^>yki^3SA^* tumors. Therefore, to test whether integrin(s) are critical for Yki-driven ISC proliferation, we down-regulated *mew* in ISCs (*esg^ts^>yki^3SA^*UAS-mew-RNAi). Strikingly, downregulation of *mew* resulted in a marked reduction in ISC proliferation (Figure 4F, G), which was more significant in the anterior midgut than in the posterior midgut. Examination of early (day 3) *esg^ts^>yki^3SA^*UAS-mew-RNAi guts revealed poor growth of ISC tumors; in particular, most of the ISCs were seen in small clusters and made up of 3 to 4 cells (Figure 4H). Further, activation of integrin alone, using a constitutively active form of the βPS integrin (49) in the ISCs (*esg^ts^>torso^D/βCyt^*), failed to trigger ISC proliferation (Figure S5), indicating that activation of the Integrin pathway is not sufficient to drive tumorigenesis in ISCs. These observations suggest that while gain of integrin signaling alone per se does not transform ISCs, it is an obligatory partner for progression of Yki-driven ISC tumors.

**Figure 4.**
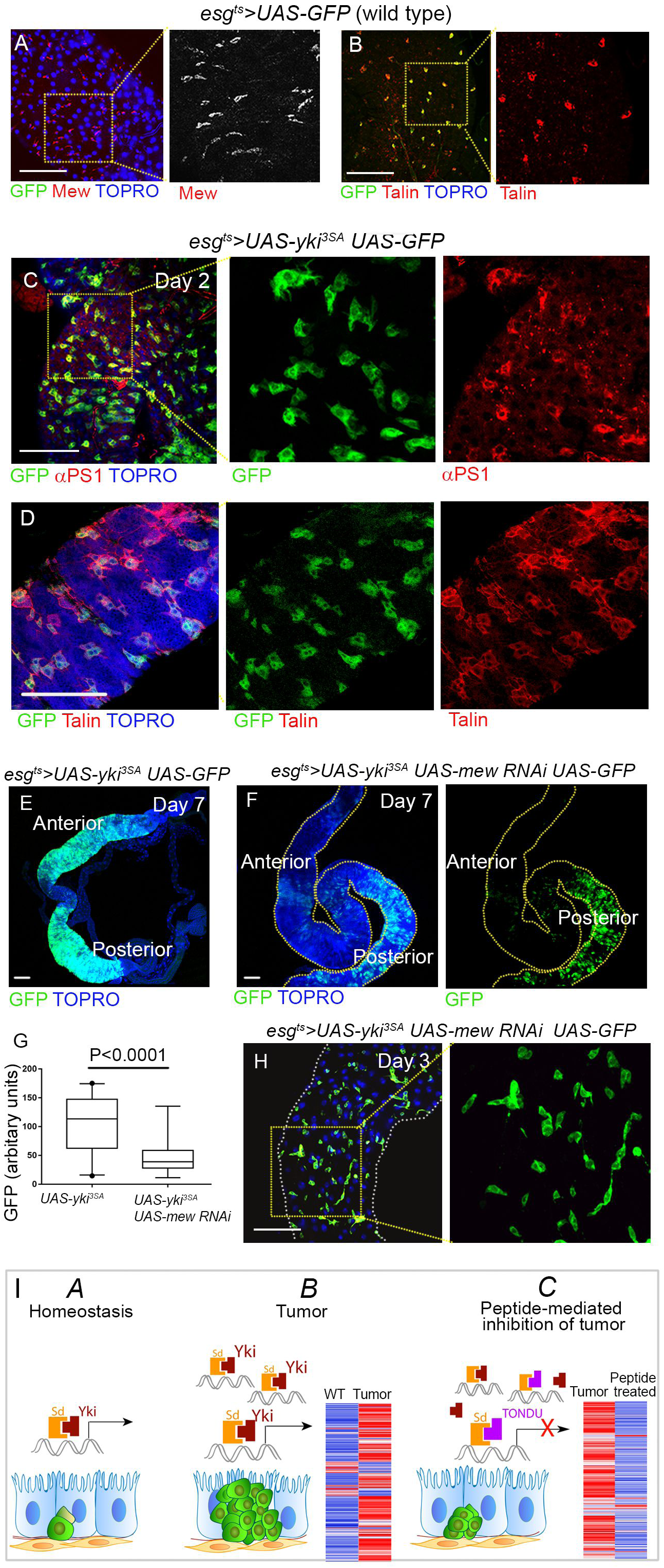
Loss of Integrin signaling inhibits growth of Yki-driven ISC tumors. (A-B) αPS1 (Mew, A) and Talin (B) staining in *esg^ts^>UAS-GFP* marked ISCs. (C-D) Overall increase in αPS1 (C) and Talin (D) in *esg^ts^>yki^3SA^* tumors. (E-F) Inhibition of Yki-driven tumors upon simultaneous down-regulation of αPS1 (*esg^ts^>yki^3SA^*UAS-mew-RNAi, n=9, F), when compared to similarly aged *esg^ts^>yki^3SA^* tumors (E). (G) Quantification of GFP from E and F. (H) Early *esg^ts^>yki^3SA^*UAS-mew-RNAi tumors (day 3) display small ISC clusters. (I) Schematic of Yki-Sd mediated transcription in wild type guts (*A*); -in Yki tumor (*B*); and in –Yki tumors in the presence of the TONDU peptide (*C*). Scale bars 100 µm.

Our observation that Yki-driven tumors could be inhibited by down-regulating integrins offers interesting therapeutic possibilities. For instance, membrane-localized integrins could be readily accessible drug targets, compared to nuclear localized YAP. Cross-species conservation of integrin signaling pathways (50) and their critical role in cancers of diverse genetic and tissue origins (13, 50) therefore presents a compelling case for integrin pathway as an alternate therapeutic target for YAP/Yki-driven cancers (51, 52). Interestingly, integrin heterodimers have been targeted either by monoclonal antibodies, such as efatucizumab and volvociximab, or by peptides such as cilengitide (53), which have proved to be effective and are currently in clinical trials. A potential caveat with this approach however is that integrin signaling is also required for wild-type ISC proliferation (41). Possibly, ISC tumors may be more sensitive to downregulation of Integrin signaling than wild type ISC, which may offer a therapeutic window for inhibitors of Integrin signaling in YAP/Yki-driven cancers.

### Concluding remarks

Advantages of low off-target activity of peptide therapy (12) are often undermined by their short half-life, poor bioavailability and uncertainties about cellular uptake. Nonetheless, targeting cellular proteins holds promise (54), as was shown earlier using the TONDU peptide (18). Our recapitulation of TONDU peptide-mediated cancer suppression (18) in ISC tumors therefore demonstrates, for the first time, that *Drosophila* could be used to screen for peptide therapeutics. Indeed, the power of genetic tractability of Drosophila, which permits generation of multiple versions of tumors of a given cell type using distinct cooperative signaling partners or transcription factors, including those seen perturbed in cancers in human (4, 6, 21, 55–57), would make such a platform versatile on several counts: scalability, genetic tractability and rapid elucidation of the mechanistic underpinning of peptide-based tumor suppression.

## MATERIALS AND METHODS

Fly stocks were obtained from the Bloomington *Drosophila* Stock Center. Antibodies were obtained from the Developmental Studies Hybridoma Bank or received as gifts from other investigators. Detailed materials and methods are provided in Supplementary Methods.

## ACKNOWLEDGEMENTS

We thank Jose F. de Celis for the antibody against Sd, Duojia Pan for HRE-Luciferase reporter vectors, Nick Brown for the UAS-*torso^D/βCyt^* fly line, and Stephanie Mohr for comments on the manuscript. We also acknowledge proteomics service by Valerian Chem, Gurgaon, India. This study was supported by the Wellcome Trust-DBT India Alliance-Early Career Fellowship (IA/E/13/1/501271) to AB.

## SUPPLEMENTARY FIGURE LEGENDS

**Figure S1.**
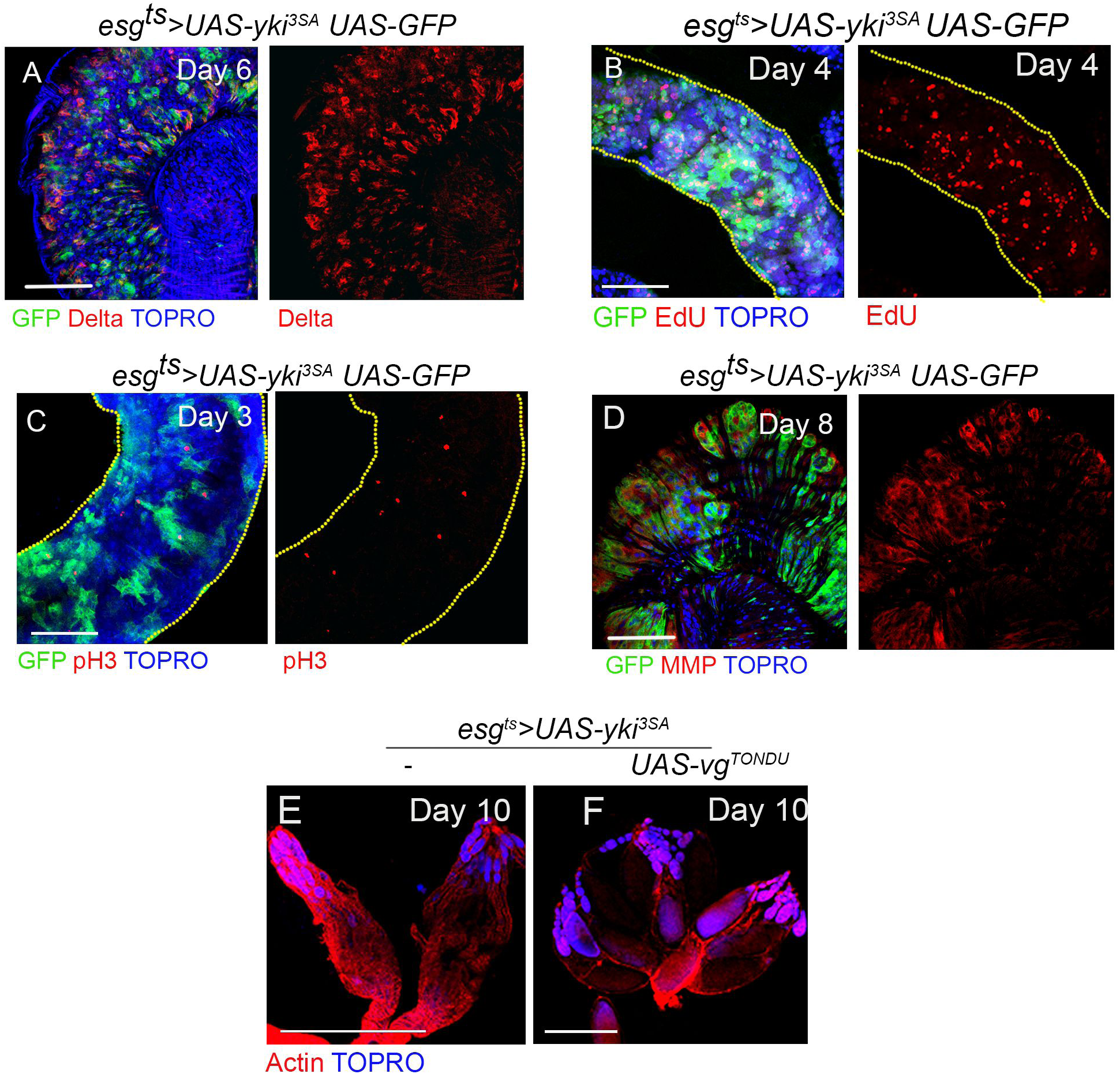
ISCs with gain of Yki display tumor phenotypes. (A-D) *esg^ts^>yki^3SA^*UAS-GFP tumors. Proliferating ISCs expressing the stem cell marker Delta (A) display an increase EdU uptake (B), increase in Phospho-Histone (C), and increase in MMP levels (D). (E) Atrophy of ovaries in *esg^ts^>yki^3SA^* flies (n=21/25). (F) Improved morphology of ovaries in *esg^ts^>yki^3SA^*UAS-vg^TONDU^ flies (n=12/25). Scale bars 100 µm.

**Figure S2.**
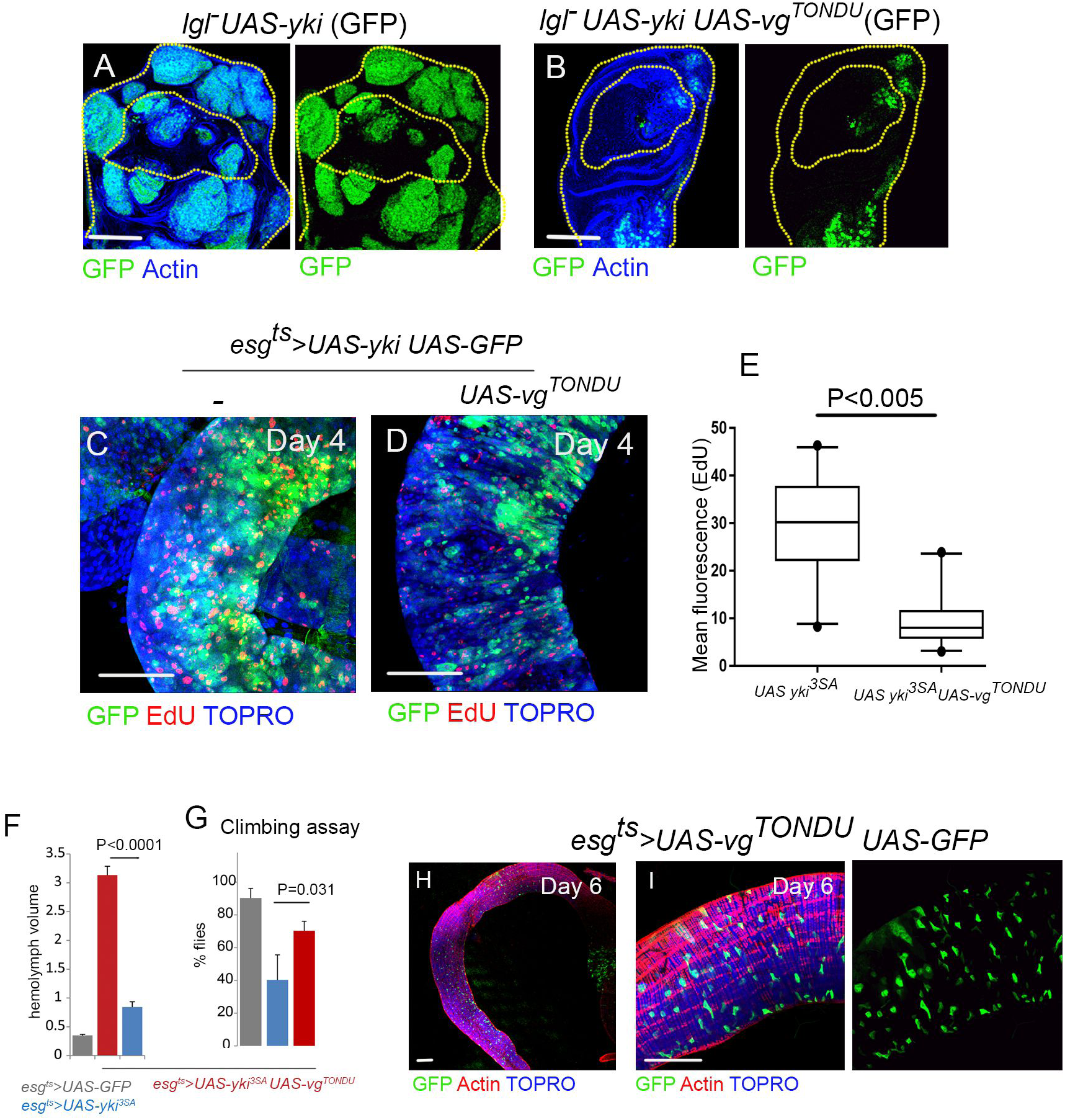
Expression of the TONDU peptide inhibits Yki-driven epithelial tumors. (A-B) Wing imaginal discs mosaic for *lgl^4^* mutant clones that express activated Yki (referred to as *lgl^4^ UAS-yki*) display tumors phenotype (A). (B) Tumor growth inhibited upon co-expression of the TONDU peptide (*lgl^4^*UAS-yki*FRT40A;UAS-vg^TONDU^*).(C-E) Decrease in the number of proliferating cells detected by EdU (red) staining in *esg^ts^>yki^3SA^*UAS-vg^TONDU^ (D), compared to *esg^ts^>yki^3SA^* tumors (C). (E) Quantification of EdU fluorescence in C and D. (F) Decrease in hemolymph content (n=25) in *esg^ts^>yki^3SA^*UAS-vg^TONDU^ flies compared to *esg^ts^>yki^3SA^* flies on Day 7. (G) TONDU-expressing *esg^ts^>yki^3SA^* flies (n=35) suppress the loss of climbing activity seen in *esg^ts^>yki^3SA^* flies. (H-I) Expression of TONDU peptide in ISCs (*esg^ts^>UAS-vg^TONDU^*) does not affect ISC numbers. Scale bars 100 µm.

**Figure S3.**
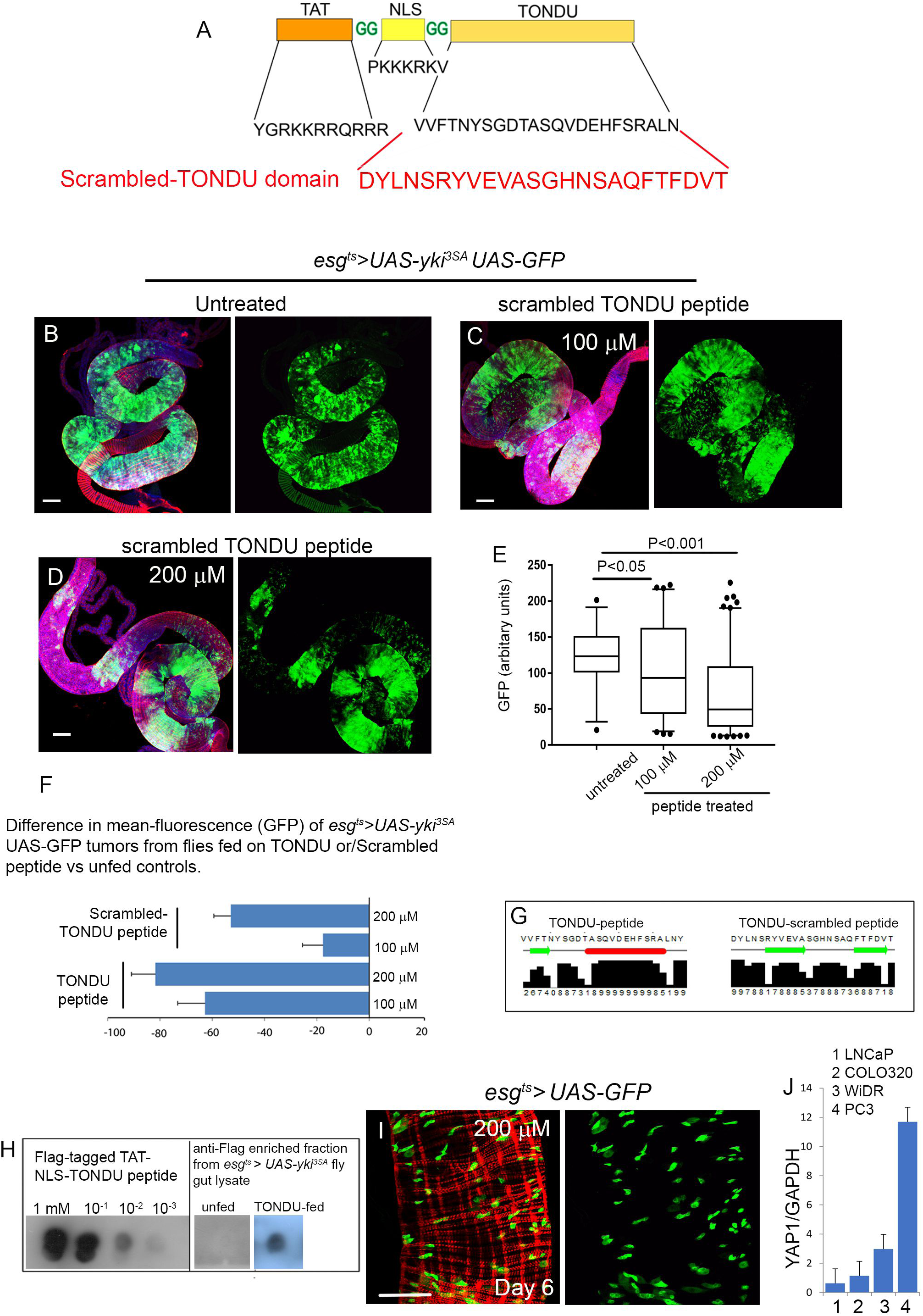
TONDU-peptide mediated inhibition of Yki-driven ISC tumors. (A) Schematic representation of the scrambled-TONDU peptide. (B-D) The scrambled-TONDU peptide displays poor growth inhibition of *esg^ts^>yki^3SA^* tumors (compare with Figure 2H and I). (E) Box plot depicting GFP quantification in *esg^ts^>yki^3SA^* tumors from flies fed on scrambled TONDU peptide. (F) Histogram displaying decrease in mean-GFP of *esg^ts^>yki^3SA^*UAS-GFP tumors from flies fed with TONDU peptide or with scrambled-TONDU peptide, when compared to unfed controls. Note that the decrease is significantly more in TONDU peptide as compared to scrambled peptide fed tumors. (G) Secondary structures of the TONDU (left) and Scrambled-TONDU (right) as predicted by JPred (http://www.compbio.dundee.ac.uk/jpred/). (H) Dot blot for FLAG-tagged TONDU peptide using anti-FLAG antibody, on native peptide (different serial dilutions); and in cell lysate (right panel) from guts (n=25) of flies fed on 200 µM of FLAG-tagged TONDU peptide and unfed flies used as control. (I) Control (*esg^ts^>UAS-GFP*) flies fed on 200 µM of TONDU peptide do not display changes in ISC numbers. (J) mRNA levels of *YAP1* in different human cancer cell lines as determined by qPCR. Scale bars 100 µm.

**Figure S4.**
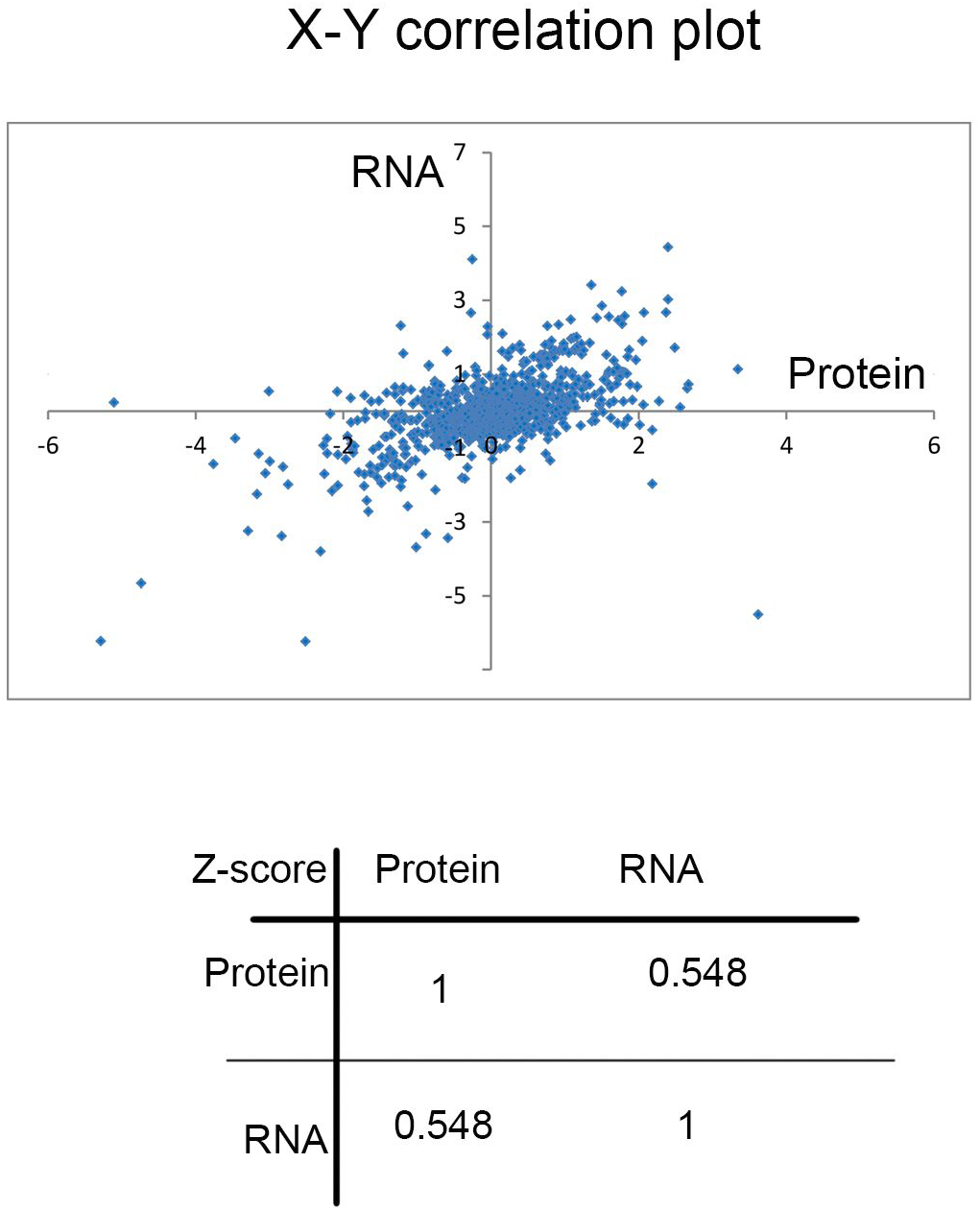
Comparison of *esg^ts^>yki^3SA^* proteome and transcriptome. x-y correlation plot displaying Z-score comparison of log_2_ fold change of genes in the proteome (current study) and transcriptome (33) of *esg^ts^>yki^3SA^* tumors.

**Figure S5.**
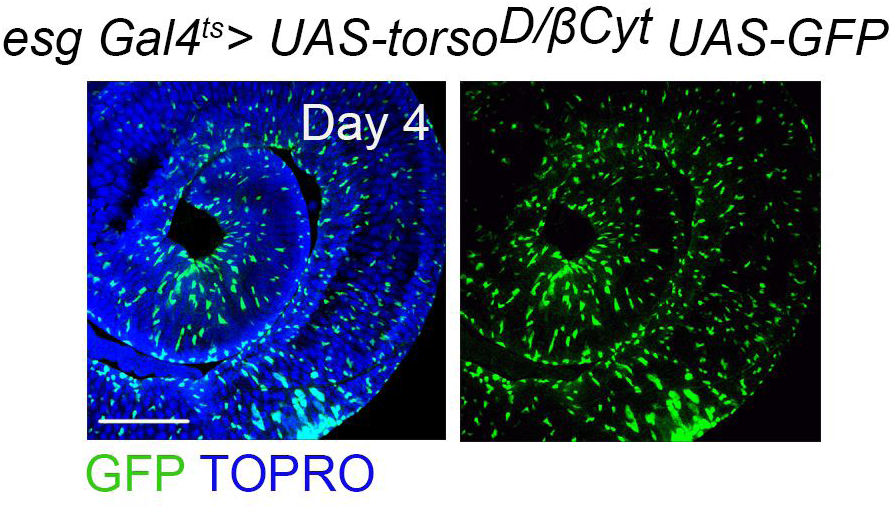
Gain of integrin signaling in *Drosophila* ISCs. Constitutive gain of integrin signaling in *esg^ts^>UAS-torso^D/βCyt^* as seen on day 4 of Gal4 activation. No aberrant proliferation of ISCs was observed. Scale bars 100 µm.

## REFERENCE

1. Vidal M, Wells S, Ryan A, & Cagan R (2005) ZD6474 suppresses oncogenic RET isoforms in a *Drosophila* model for type 2 multiple endocrine neoplasia syndromes and papillary thyroid carcinoma. Cancer Res 65(9):3538–3541.

2. Dar AC, Das TK, Shokat KM, & Cagan RL (2012) Chemical genetic discovery of targets and anti-targets for cancer polypharmacology. Nature 486(7401):80–84.

3. Willoughby LF, et al. (2013) An in vivo large-scale chemical screening platform using *Drosophila* for anti-cancer drug discovery. Dis Model Mech 6(2):521–529.

4. Levine BD & Cagan RL (2016) Drosophila Lung Cancer Models Identify Trametinib plus Statin as Candidate Therapeutic. Cell Rep 14(6):1477–1487.

5. Markstein M, et al. (2014) Systematic screen of chemotherapeutics in *Drosophila* stem cell tumors. Proc Natl Acad Sci U S A 111(12):4530–4535.

6. Bangi E, Murgia C, Teague AG, Sansom OJ, & Cagan RL (2016) Functional exploration of colorectal cancer genomes using Drosophila. Nat Commun 7:13615.

7. Darnell JE, Jr. (2002) Transcription factors as targets for cancer therapy. Nat Rev Cancer 2(10):740–749.

8. Lambert M, Jambon S, Depauw S, & David-Cordonnier MH (2018) Targeting Transcription Factors for Cancer Treatment. Molecules 23(6).

9. Lau JL & Dunn MK (2018) Therapeutic peptides: Historical perspectives, current development trends, and future directions. Bioorg Med Chem 26(10):2700–2707.

10. Drucker DJ (2019) Advances in oral peptide therapeutics. Nat Rev Drug Discov.

11. Ley K, Rivera-Nieves J, Sandborn WJ, & Shattil S (2016) Integrin-based therapeutics: biological basis, clinical use and new drugs. Nat Rev Drug Discov 15(3):173–183.

12. Furet P, et al. (2019) Structure-based design of potent linear peptide inhibitors of the YAP-TEAD protein-protein interaction derived from the YAP omega-loop sequence. Bioorg Med Chem Lett 29(16):2316–2319.

13. Zanconato F, Cordenonsi M, & Piccolo S (2019) YAP and TAZ: a signalling hub of the tumour microenvironment. Nat Rev Cancer 19(8):454–464.

14. Huh HD, Kim DH, Jeong HS, & Park HW (2019) Regulation of TEAD Transcription Factors in Cancer Biology. Cells 8(6).

15. Pobbati AV, Chan SW, Lee I, Song H, & Hong W (2012) Structural and functional similarity between the Vgll1-TEAD and the YAP-TEAD complexes. Structure 20(7):1135–1140.

16. Zhang W, et al. (2014) VGLL4 functions as a new tumor suppressor in lung cancer by negatively regulating the YAP-TEAD transcriptional complex. Cell Res 24(3):331–343.

17. Zhang Y, et al. (2017) VGLL4 Selectively Represses YAP-Dependent Gene Induction and Tumorigenic Phenotypes in Breast Cancer. Sci Rep 7(1):6190.

18. Jiao S, et al. (2014) A peptide mimicking VGLL4 function acts as a YAP antagonist therapy against gastric cancer. Cancer Cell 25(2):166–180.

19. Huang J, Wu S, Barrera J, Matthews K, & Pan D (2005) The Hippo signaling pathway coordinately regulates cell proliferation and apoptosis by inactivating Yorkie, the *Drosophila* Homolog of YAP. Cell 122(3):421–434.

20. Cho E, et al. (2006) Delineation of a Fat tumor suppressor pathway. Nat Genet 38(10):1142–1150.

21. Khan SJ, et al. (2013) Epithelial neoplasia in *Drosophila* entails switch to primitive cell states. Proc Natl Acad Sci U S A 110(24):E2163–2172.

22. Menendez J, Perez-Garijo A, Calleja M, & Morata G (2010) A tumor-suppressing mechanism in *Drosophila* involving cell competition and the Hippo pathway. Proc Natl Acad Sci U S A 107(33):14651–14656.

23. Wu S, Liu Y, Zheng Y, Dong J, & Pan D (2008) The TEAD/TEF family protein Scalloped mediates transcriptional output of the Hippo growth-regulatory pathway. Dev Cell 14(3):388–398.

24. Zhang L, et al. (2008) The TEAD/TEF family of transcription factor Scalloped mediates Hippo signaling in organ size control. Dev Cell 14(3):377–387.

25. Halder G, et al. (1998) The Vestigial and Scalloped proteins act together to directly regulate wing-specific gene expression in Drosophila. Genes Dev 12(24):3900–3909.

26. Simmonds AJ, et al. (1998) Molecular interactions between Vestigial and Scalloped promote wing formation in Drosophila. Genes Dev 12(24):3815–3820.

27. Vaudin P, Delanoue R, Davidson I, Silber J, & Zider A (1999) TONDU (TDU), a novel human protein related to the product of vestigial (vg) gene of *Drosophila* melanogaster interacts with vertebrate TEF factors and substitutes for Vg function in wing formation. Development 126(21):4807–4816.

28. Karpowicz P, Perez J, & Perrimon N (2010) The Hippo tumor suppressor pathway regulates intestinal stem cell regeneration. Development 137(24):4135–4145.

29. Staley BK & Irvine KD (2010) Warts and Yorkie mediate intestinal regeneration by influencing stem cell proliferation. Curr Biol 20(17):1580–1587.

30. Ren F, et al. (2010) Hippo signaling regulates *Drosophila* intestine stem cell proliferation through multiple pathways. Proc Natl Acad Sci U S A 107(49):21064–21069.

31. Jin Y, et al. (2013) Brahma is essential for *Drosophila* intestinal stem cell proliferation and regulated by Hippo signaling. Elife 2:e00999.

32. Kwon Y, et al. (2015) Systemic organ wasting induced by localized expression of the secreted insulin/IGF antagonist ImpL2. Dev Cell 33(1):36–46.

33. Song W, et al. (2019) Tumor-Derived Ligands Trigger Tumor Growth and Host Wasting via Differential MEK Activation. Dev Cell 48(2):277–286 e276.

34. Guo Z, Lucchetta E, Rafel N, & Ohlstein B (2016) Maintenance of the adult *Drosophila* intestine: all roads lead to homeostasis. Curr Opin Genet Dev 40:81–86.

35. Figueroa-Clarevega A & Bilder D (2015) Malignant *Drosophila* tumors interrupt insulin signaling to induce cachexia-like wasting. Dev Cell 33(1):47–55.

36. Oh H & Irvine KD (2008) In vivo regulation of Yorkie phosphorylation and localization. Development 135(6):1081–1088.

37. Wadia JS & Dowdy SF (2005) Transmembrane delivery of protein and peptide drugs by TAT-mediated transduction in the treatment of cancer. Adv Drug Deliv Rev 57(4):579–596.

38. Szklarczyk D, et al. (2019) STRING v11: protein-protein association networks with increased coverage, supporting functional discovery in genome-wide experimental datasets. Nucleic Acids Res 47(D1):D607–D613.

39. Mulligan KA, et al. (2012) Secreted Wingless-interacting molecule (Swim) promotes long-range signaling by maintaining Wingless solubility. Proc Natl Acad Sci U S A 109(2):370–377.

40. Kwon Y, et al. (2013) The Hippo signaling pathway interactome. Science 342(6159):737–740.

41. Chen J, Sayadian AC, Lowe N, Lovegrove HE, & St Johnston D (2018) An alternative mode of epithelial polarity in the *Drosophila* midgut. PLoS Biol 16(10):e3000041.

42. Lin G, et al. (2013) Integrin signaling is required for maintenance and proliferation of intestinal stem cells in Drosophila. Dev Biol 377(1):177–187.

43. Oh H, et al. (2013) Genome-wide association of Yorkie with chromatin and chromatin-remodeling complexes. Cell Rep 3(2):309–318.

44. Nagaraj R, et al. (2012) Control of mitochondrial structure and function by the Yorkie/YAP oncogenic pathway. Genes Dev 26(18):2027–2037.

45. Guilgur LG, et al. (2014) Requirement for highly efficient pre-mRNA splicing during *Drosophila* early embryonic development. Elife 3:e02181.

46. Biswas S, Lunec J, & Bartlett K (2012) Non-glucose metabolism in cancer cells--is it all in the fat? Cancer Metastasis Rev 31(3-4):689–698.

47. Koundouros N & Poulogiannis G (2020) Reprogramming of fatty acid metabolism in cancer. Br J Cancer 122(1):4–22.

48. Reis R, et al. (2009) Dynein and mast/orbit/CLASP have antagonistic roles in regulating kinetochore-microtubule plus-end dynamics. J Cell Sci 122(Pt 14):2543–2553.

49. Martin-Bermudo MD, Alvarez-Garcia I, & Brown NH (1999) Migration of the *Drosophila* primordial midgut cells requires coordination of diverse PS integrin functions. Development 126(22):5161–5169.

50. Cooper J & Giancotti FG (2019) Integrin Signaling in Cancer: Mechanotransduction, Stemness, Epithelial Plasticity, and Therapeutic Resistance. Cancer Cell 35(3):347–367.

51. Arun AS, Tepper CG, & Lam KS (2018) Identification of integrin drug targets for 17 solid tumor types. Oncotarget 9(53):30146–30162.

52. Arosio D, Manzoni L, Corno C, & Perego P (2017) Integrin-Targeted Peptide- and Peptidomimetic-Drug Conjugates for the Treatment of Tumors. Recent Pat Anticancer Drug Discov 12(2):148–168.

53. Raab-Westphal S, Marshall JF, & Goodman SL (2017) Integrins as Therapeutic Targets: Successes and Cancers. Cancers (Basel) 9(9).

54. Chang YS, et al. (2013) Stapled alpha-helical peptide drug development: a potent dual inhibitor of MDM2 and MDMX for p53-dependent cancer therapy. Proc Natl Acad Sci U S A 110(36):E3445–3454.

55. Gupta RP, Bajpai A, & Sinha P (2017) Selector genes display tumor cooperation and inhibition in *Drosophila* epithelium in a developmental context-dependent manner. Biology open 6(11):1581–1591.

56. Bajpai A & Sinha P (2019) Hh signaling from de novo organizers drive lgl neoplasia in *Drosophila* epithelium. Dev Biol.

57. Das TK & Cagan RL (2017) KIF5B-RET Oncoprotein Signals through a Multi-kinase Signaling Hub. Cell Rep 20(10):2368–2383.

58. Mi H, Guo N, Kejariwal A, & Thomas PD (2007) PANTHER version 6: protein sequence and function evolution data with expanded representation of biological pathways. Nucleic Acids Res 35(Database issue):D247–252.

