## Supplementary Information for "A *Drosophila* model of oral peptide therapeutics for adult Intestinal Stem Cell tumors"

**Table S1:** Protein abundance in $esg^{ts}$>$yki^{3SA}$ driven ISC tumors on day 7 and day 1 of tumor induction. (Provided as an excel sheet).

**Table S2:** Proteins with significant fold change ($\geq \log_2 2$, $p$<0.05) in day 7 versus day 1 of $esg^{ts}$>$yki^{3SA}$ driven ISC tumors.

**Table S3:** Interacting partners belonging to the Hippo pathway for proteins with $\geq \log_2 2$ fold change in $esg^{ts}$>$yki^{3SA}$ driven ISC tumors.

**Table S4:** Gene Ontology (GO) analysis of enriched proteins in $esg^{ts}$>$yki^{3SA}$ driven ISC tumors.

**Table S5**: Status of integrin pathway members in $esg^{ts}$>$yki^{3SA}$ driven ISC tumors.

**Table S6:** Putative Yki-Sd binding sites in regulatory regions of genes with $>\log_2 2$ fold change in protein levels $esg^{ts}$>$yki^{3SA}$ driven ISC tumors.

**Table S7:** Fold change in protein levels in TONDU peptide-treated $esg^{ts}$>$yki^{3SA}$ driven ISC tumors.

**Table S2:** Proteins with significant fold change (≥$\log_2 2$, $p$<0.05) in day 7 versus day 1 of $esg^{ts}$>$yki^{3SA}$ driven ISC tumors.

| Gene ID | Gene Symbol | Protein FDR Confidence: Combined | Master | UniProt Accession | Median *(Day 1)* | Median *(Day 7)* | Abundance Ratio: Day 7/ Day 1 | Abundance Ratio (log2) | T-Test (P value) | # amino acis | MW [kDa] | # Peptides (SequestHT) | Chromosome | Found in Sample Group: (Day 1) | Found in Sample Group: (Day 7) |
|---|---|---|---|---|---|---|---|---|---|---|---|---|---|---|---|
| **FBgn0261276** | *Opa1* | High | None | F0JAH2 | 74046.703 | 297394.711 | 4.016 | **2.006** | **0.0048** | 453 | 51.2 | 2 | 2R | Peak Found | High |
| **FBgn0000562** | *egl* | Medium | Master Protein Candidate | Q9W1K4 | 14056.321 | 57460.557 | 4.088 | **2.031** | **0.0430** | 1004 | 112.1 | 1 | 2R | Peak Found | High |
| **FBgn0260442** | *rhea* | High | Master Protein | Q960C2 | 67243.947 | 276212.477 | 4.108 | **2.038** | **0.0264** | 1601 | 171.2 | 26 | 3L | High | High |
| **FBgn0035498** | *Fit1* | High | Master Protein | Q9VZI3 | 394737.461 | 1640060.666 | 4.155 | **2.055** | **0.0276** | 708 | 80.4 | 6 | 3L | High | High |
| **FBgn0035793** | *CG7546* | High | Master Protein Candidate | M9PBU3 | 91147.313 | 390343.234 | 4.283 | **2.098** | **0.0087** | 1179 | 125.7 | 2 | 3L | High | High |
| **FBgn0030955** | *CG6891* | High | Master Protein Candidate | Q8MQZ6 | 100332.746 | 448638.875 | 4.472 | **2.161** | **0.0323** | 269 | 30.2 | 1 | X | High | High |
| **FBgn0013437** | *copia* | High | Master Protein | P04146 | 581703.875 | 2664281.188 | 4.580 | **2.195** | **0.0138** | 1409 | 162.7 | 3 | | High | High |
| **FBgn0265991** | *Zasp52* | High | None | G3JX29 | 147506.977 | 678663.543 | 4.601 | **2.202** | **0.0366** | 651 | 70.9 | 3 | 2R | High | High |
| **FBgn0015379** | *dod* | High | Master Protein | P54353 | 10240.124 | 94620.563 | 9.240 | **3.208** | **0.0451** | 166 | 18.4 | 1 | X | Peak Found | High |
| **FBgn0000667** | *Actn* | High | Master Protein Candidate | M9MS06 | 60172.696 | 284884.955 | 4.734 | **2.243** | **0.0489** | 895 | 103.8 | 4 | X | Peak Found | High |
| **FBgn0010434** | *cora* | High | Master Protein Candidate | A0A0B4 LFX4 | 469318.387 | 2248560.410 | 4.791 | **2.260** | **0.0047** | 1600 | 173.8 | 6 | 2R | High | High |
| **FBgn0014020** | *Rho1* | High | Master Protein | P48148 | 491985.963 | 2396139.059 | 4.870 | **2.284** | **0.0177** | 192 | 21.7 | 6 | 2R | Peak Found | High |
| **FBgn0030052** | *CG12065* | High | None | Q8MRM6 | 279747.601 | 1375997.625 | 4.919 | **2.298** | **0.0460** | 641 | 71.1 | 5 | X | Peak Found | High |
| **FBgn0053470** | *CG33470; IM10; IMPPP* | High | Master Protein | Q8ML70 | 3874.029 | 38433.805 | 9.921 | **3.310** | **0.0377** | 257 | 28 | 1 | 2R | Peak Found | High |
| **FBgn0035499** | *Chd64* | High | None | M9PE30 | 6518473.696 | 32604473.558 | 5.002 | **2.322** | **0.0003** | 175 | 19.4 | 25 | 3L | High | High |
| **FBgn0025352** | *Mtpβ* | High | Master Protein | O77466 | 60337.865 | 605581.250 | 10.037 | **3.327** | **0.0340** | 469 | 50.6 | 1 | 2R | Peak Found | High |
| **FBgn0086346** | *ALiX* | High | Master Protein | Q9VB05 | 77987.759 | 397127.230 | 5.092 | **2.348** | **0.0169** | 836 | 92.5 | 2 | 3R | Peak Found | High |
| **FBgn0260442** | *rhea* | High | None | M9NDM3 | 1072195.987 | 5647432.310 | 5.267 | **2.397** | **0.0350** | 2169 | 235.1 | 31 | 3L | High | High |
| **FBgn0004397** | *Vinc* | High | Master Protein | X2JAB9 | 741152.635 | 3905646.674 | 5.270 | **2.398** | **0.0224** | 961 | 106.2 | 12 | X | High | High |
| **FBgn0021760** | *chb* | High | Master Protein | Q9NBD7 | 31028.359 | 169383.656 | 5.459 | **2.449** | **0.0329** | 1491 | 165.5 | 2 | 3L | Peak Found | High |
| **FBgn0262567** | *CG43107* | High | Master Protein | D0IQC0 | 9607.135 | 107383.602 | 11.177 | **3.483** | **0.0174** | 63 | 6.8 | 1 | 2R | Peak Found | Peak Found |
| **FBgn0013733** | *shot* | High | Master Protein | A1Z9J3 | 28039.734 | 350253.043 | 12.491 | **3.643** | **0.0138** | 8805 | 988.9 | 2 | 2R | Peak Found | High |
| **FBgn0033446** | *CG1648* | High | Master Protein | Q7K2P3 | 775279.577 | 5030252.721 | 6.488 | **2.698** | **0.0201** | 230 | 23.8 | 18 | 2R | High | High |
| **FBgn0262735** | *Imp* | High | Master Protein | Q0KHU2 | 88808.606 | 580638.375 | 6.538 | **2.709** | **0.0201** | 631 | 69.4 | 3 | X | Peak Found | High |
| **FBgn0022709** | *Adk1* | High | Master Protein Candidate | Q9VTV3 | 17311.554 | 113919.848 | 6.581 | **2.718** | **0.0072** | 201 | 21.9 | 1 | 3L | Peak Found | High |
| **FBgn0086906** | *sls* | High | None | R4UAY6 | 23315.416 | 308855.279 | 13.247 | **3.728** | **0.0122** | 662 | 74.6 | 2 | 3L | High | High |
| **FBgn0010909** | *msn* | Medium | None | Q7KV90 | 7461.706 | 102635.387 | 13.755 | **3.782** | **0.0121** | 1200 | 130.3 | 1 | 3L | Peak Found | High |

| | | | | | | | | | | | | | | | |
|---|---|---|---|---|---|---|---|---|---|---|---|---|---|---|---|
| **FBgn0036580** | *PDCD-5* | High | Master Protein | Q9VUZ8 | 11643.499 | 169815.949 | 14.585 | **3.866** | **0.0111** | 133 | 15.1 | 2 | 3L | Peak Found | Peak Found |
| **FBgn0051901** | *Mur29B* | Medium | Master Protein Candidate | Q8MS63 | 22220.121 | 166646.781 | 7.500 | **2.907** | **0.0366** | 339 | 35.7 | 1 | 2L | High | High |
| **FBgn0265434** | *zip* | High | None | A0A0B4JD95 | 3745790.734 | 28852286.631 | 7.703 | **2.945** | **0.0010** | 1964 | 226.6 | 67 | 2R | High | High |
| **FBgn0005666** | *bent* | High | None | O76281 | 284599.331 | 2219016.392 | 7.797 | **2.963** | **0.0385** | 6658 | 743 | 15 | 4 | Peak Found | High |
| **FBgn0262716** | *Arp3; Arp66B* | High | Master Protein | P32392 | 91762.304 | 725246.529 | 7.904 | **2.982** | **0.0397** | 418 | 47 | 4 | 3L | High | High |
| **FBgn0053113** | *Rtnl1* | High | Master Protein Candidate | Q9VMV9 | 594076.854 | 5048772.010 | 8.499 | **3.087** | **0.0238** | 595 | 63.9 | 9 | 2L | High | High |
| **FBgn0085282** | *CG34253* | High | Master Protein | A8JNV2 | 26812.808 | 228363.336 | 8.517 | **3.090** | **0.0221** | 112 | 12.5 | 1 | 3L | Peak Found | High |
| **FBgn0265434** | *zip* | High | None | J7JVR0 | 1687952.657 | 14541350.003 | 8.615 | **3.107** | **0.0003** | 1425 | 164.4 | 39 | 2R | High | High |
| **FBgn0016726** | *RpL29* | High | Master Protein | B7FNL1 | 732637.288 | 6371885.220 | 8.697 | **3.121** | **0.0452** | 85 | 10 | 6 | 2R | High | High |
| **FBgn0029766** | *CG15784* | High | Master Protein | Q9W4C1 | 151556.702 | 1363172.164 | 8.994 | **3.169** | **0.0261** | 554 | 62.3 | 5 | X | High | High |
| **FBgn0038774** | *CG5023* | High | Master Protein | Q9I7J0 | 580277.645 | 5744084.078 | 9.899 | **3.307** | **0.0314** | 169 | 19.1 | 9 | 3R | High | High |
| **FBgn0001217** | *Hsc70-2* | High | Master Protein | P11146 | 34573.184 | 694860.797 | 20.098 | **4.329** | **0.0135** | 633 | 69.7 | 3 | 3R | Peak Found | High |
| **FBgn0051363** | *CG31363* | High | Master Protein | B5RJ67 | 30460.900 | 612434.938 | 20.106 | **4.330** | **0.0112** | 230 | 24.4 | 1 | 3R | Peak Found | High |
| **FBgn0000043** | *Act42A* | High | Master Protein | P02572 | 13317.133 | 326000.129 | 24.480 | **4.614** | **0.0155** | 376 | 41.8 | 37 | 2R | Peak Found | High |
| **FBgn0004227** | *nonA* | High | Master Protein | Q8IR16 | 51621.380 | 664389.951 | 12.870 | **3.686** | **0.0092** | 742 | 81.9 | 5 | X | High | High |
| **FBgn0011225** | *jar* | High | Master Protein | Q01989 | 32407.746 | 421589.398 | 13.009 | **3.701** | **0.0170** | 1253 | 143.2 | 6 | 3R | Peak Found | High |
| **FBgn0034709** | *CG3074; Swim* | High | Master Protein | Q7JWQ7 | 582800.990 | 7699009.721 | 13.210 | **3.724** | **0.0014** | 431 | 48.8 | 11 | 2R | High | High |
| **FBgn0031436** | *CG3214; ND-B17.2* | High | Master Protein | Q9VQD7 | 13005.392 | 404752.090 | 31.122 | **4.960** | **0.0032** | 142 | 16.8 | 2 | 2L | High | High |
| **FBgn0000639** | *Fbp1* | High | Master Protein Candidate | M9PFK6 | 2941545.182 | 75592.226 | 0.026 | **-5.282** | **0.0184** | 1027 | 119.4 | 15 | 3L | High | Peak Found |
| **FBgn0033297** | *Mal-A8* | High | Master Protein Candidate | H5V882 | 145483.008 | 4449.611 | 0.031 | **-5.031** | **0.0020** | 597 | 68.1 | 1 | 2R | High | Peak Found |
| **FBgn0010497** | *dmGlut; l(2)01810* | Medium | Master Protein Candidate | Q95R95 | 217644.070 | 7972.730 | 0.037 | **-4.771** | **0.0143** | 165 | 17.7 | 1 | 2L | High | Peak Found |
| **FBgn0004426** | *LysC* | High | None | P83971 | 1020142.977 | 101441.592 | 0.099 | **-3.330** | **0.0171** | 140 | 15.6 | 2 | 3L | High | Peak Found |
| **FBgn0031141** | *CG1304* | High | None | Q9VRD1 | 1379758.828 | 251684.488 | 0.182 | **-2.455** | **0.0240** | 260 | 27.8 | 1 | X | High | High |
| **FBgn0004427** | *LysD* | High | Master Protein Candidate | P83972 | 2366383.414 | 523926.295 | 0.221 | **-2.175** | **0.0126** | 140 | 15.6 | 3 | 3L | High | High |
| **FBgn0004425** | *LysB* | High | Master Protein Candidate | Q08694 | 2366383.414 | 523926.295 | 0.221 | **-2.175** | **0.0126** | 140 | 15.6 | 3 | 3L | High | High |
| **FBgn0004428** | *LysE* | High | Master Protein | P37159 | 2366383.414 | 523926.295 | 0.221 | **-2.175** | **0.0126** | 140 | 15.5 | 3 | 3L | High | High |
| **FBgn0034662** | *CG13492* | High | Master Protein | Q8MLU9 | 1715613.215 | 393704.303 | 0.229 | **-2.124** | **0.0428** | 2979 | 321.1 | 12 | 2R | High | High |
| **FBgn0040349** | *CG3699* | High | Master Protein Candidate | Q9U1L2 | 6207241.123 | 1490634.446 | 0.240 | **-2.058** | **0.0024** | 251 | 26 | 12 | X | High | High |

**Table S3:** Interacting partners belonging to the Hippo pathway for proteins with $\geq \log_2 2$ fold change in *esg*$^{ts}$>*yki*$^{3SA}$ driven ISC tumors.

.

| Protein | Fold change ($\log_2$ fold) | T Test P value | | Uniprot | $Hippo protein-protein interaction network | |
|---|---|---|---|---|---|---|
| | | | | | Protein-and-Hippo pathway member | Score |
| Cora | 2.2604 | 0.0047 | High | A0A0B4LFX4 | Cora ←→Ft | 1* |
| Mtb-β | 3.3272 | 0.0340 | Peak Found | O77466 | Mtb-β ←→Ft | 0.99* |
| Msn | 3.7819 | 0.0121 | Peak Found | Q7KV90 | Msn ←→Ft | 1* |
| nonA | 3.6860 | 0.0092 | High | Q8IR16 | nonA ←→Ex | 0.82* |
| Jar | 3.7014 | 0.0170 | Peak Found | Q01989 | Jar ←→Wts | 0.89* |
| Talin | 2.0383 | 0.0264 | High | Q960C2 | Talin ←→Yki | 0.33 |
| Vinc | 2.2184 | 0.0188 | High | Q24584 | Vinc ←→Ft | 0.36 |
| Chd64 | 2.3225 | 0.0003 | High | M9PE30 | Chd64 ←→Yki | 0.01 |
| Mtb-β | 3.3272 | 0.0340 | Peak Found | O77466 | Mtb-β ←→Wts | 0.1 |
| Arp3 | 2.9825 | 0.0397 | High | P32392 | Arp3 ←→Wts | 0.21 |
| Rtnl1 | 3.0872 | 0.0238 | High | Q9VMV9 | Rtnl1 ←→Ft | 0.38 |
| nonA | 3.6860 | 0.0092 | High | Q8IR16 | nonA ←→Wts | 0.28 |

$Kwon *et al.* (1). *Statistically significant intections

**Table S4:** Gene Ontology (GO) analysis of enriched proteins in *esg*$^{ts}$>*yki*$^{3SA}$ driven ISC tumors.

| MOLECULAR FUNCTION | | $GO classes | Gene Numbers | % representation | % representation |
|---|---|---|---|---|---|
| | 1 | binding (GO:0005488) | 7 | 19.40% | 29.20% |
| | 2 | structural molecule activity (GO:0005198) | 6 | 16.70% | 25.00% |
| | 3 | molecular function regulator (GO:0098772) | 1 | 2.80% | 4.20% |
| | 4 | catalytic activity (GO:0003824) | 10 | 27.80% | 41.70% |
| **BIOLOGICAL PROCESS** | | | | | |
| | 1 | response to stimulus (GO:0050896) | 2 | 5.60% | 7.40% |
| | 2 | cellular process (GO:0009987) | 11 | 30.60% | 40.70% |
| | 3 | multicellular organismal process (GO:0032501) | 1 | 2.80% | 3.70% |
| | 4 | metabolic process (GO:0008152) | 6 | 16.70% | 22.20% |
| | 5 | biological regulation (GO:0065007) | 2 | 5.60% | 7.40% |
| | 6 | localization (GO:0051179) | 4 | 11.10% | 14.80% |
| | 7 | biological adhesion (GO:0022610) | 1 | 2.80% | 3.70% |
| **CELLULAR COMPONENT** | | | | | |
| | 1 | organelle (GO:0043226) | 9 | 25.00% | 45.00% |
| | 2 | extracellular region (GO:0005576) | 1 | 2.80% | 5.00% |
| | 3 | cell (GO:0005623) | 10 | 27.80% | 50.00% |
| **PROTEIN CLASSES** | | | | | |
| | 1 | transmembrane receptor regulatory/adaptor protein (PC00226) | 1 | 2.80% | 4.20% |
| | 2 | hydrolase (PC00121) | 5 | 13.90% | 20.80% |
| | 3 | cell junction protein (PC00070) | 1 | 2.80% | 4.20% |

| | | | | | |
|---|---|---|---|---|---|
| | 4 | enzyme modulator (PC00095) | 6 | 16.70% | 25.00% |
| | 5 | nucleic acid binding (PC00171) | 3 | 8.30% | 12.50% |
| | 6 | transferase (PC00220) | 1 | 2.80% | 4.20% |
| | 7 | receptor (PC00197) | 1 | 2.80% | 4.20% |
| | 8 | cytoskeletal protein (PC00085) | 5 | 13.90% | 20.80% |
| | 9 | structural protein (PC00211) | 1 | 2.80% | 4.20% |
| **PATHWAYS** | | | | | |
| | 1 | Gonadotropin-releasing hormone receptor pathway (P06664) | 1 | 2.80% | 4.30% |
| | 2 | Cadherin signaling pathway (P00012) | 1 | 2.80% | 4.30% |
| | 3 | De novo purine biosynthesis (P02738) | 1 | 2.80% | 4.30% |
| | 4 | Axon guidance mediated by Slit/Robo (P00008) | 1 | 2.80% | 4.30% |
| | 5 | Apoptosis signaling pathway (P00006) | 1 | 2.80% | 4.30% |
| | 6 | Integrin signalling pathway (P00034) | 5 | 13.90% | 21.70% |
| | 7 | Angiogenesis (P00005) | 1 | 2.80% | 4.30% |
| | 8 | Alzheimer disease-presenilin pathway (P00004) | 1 | 2.80% | 4.30% |
| | 9 | Inflammation mediated by chemokine and cytokine signaling pathway (P00031) | 2 | 5.60% | 8.70% |
| | 10 | Huntington disease (P00029) | 1 | 2.80% | 4.30% |
| | 11 | Parkinson disease (P00049) | 1 | 2.80% | 4.30% |
| | 12 | Ras Pathway (P04393) | 1 | 2.80% | 4.30% |
| | 13 | Cytoskeletal regulation by Rho GTPase (P00016) | 3 | 8.30% | 13.00% |
| | 14 | Nicotinic acetylcholine receptor signaling pathway (P00044) | 3 | 8.30% | 13.00% |

[$]GO enrichment was determined using PANTHER (http://www.pantherdb.org/).

**Table S5:** Status of integrin pathway members in $esg^{ts}>yki^{3SA}$ driven ISC tumors.

| Gene ID | Pathway member | Fly base ID | **Proteomics** ($log_2$ fold). Current study | [$]**RNAseq** ($log_2$ fold) |
|---|---|---|---|---|
| *scb/aPS3* | Integrin receptor | FBgn0003328 | 2.6069 (UP, P=0.270) | 3.2797 (UP) |
| *rhea* | Adaptor | FBgn0260442 | 2.3970 (UP, P=0.035) | 2.4646 (UP) |
| *Ilk* | Kinase and Scaffold protein | FBgn0028427 | 2.0724 (UP, P=0.056) | 1.6871 (UP) |
| *Pax* | Scaffold protein | FBgn0041789 | 6.0551 (UP, P=0.051) | 1.6507 (UP) |
| *Vinc* | Scaffold protein | FBgn0004397 | 2.3977 (UP, P=0.022) | 2.8706 (UP) |
| *vkg* | Basemen Membrane | FBgn0016075 | 1.8255 (UP, P=0.014) | 2.5527 (UP) |
| *Rho1* | GTPase | FBgn0014020 | 2.2840 (UP, P=0.018) | 2.3794 (UP) |
| *Act42A* | Cytoskeleton | FBgn0000043 | 4.6013 (UP, P=0.1934) | 5.1188 (UP) |
| *mew/alpha-PS1* | Integrin receptor | FBgn0004456 | not found | 3.3184 (UP) |
| *mys/betaPS1* | Integrin receptor | FBgn0004657 | not found | 3.7469 (UP) |
| *if/alphaPS2,* | Integrin receptor | FBgn0001250 | not found | 0.3507 |
| *Itgbn/Itgbetanu* | Integrin receptor | FBgn0010395 | not found | 0.1434 |
| *LanA* | Ligand | FBgn0002526 | not found | 3.0021 (UP) |
| *wb/LanA1* | Ligand | FBgn0261563 | not found | 2.0211 (UP) |
| *LanB1* | Ligand | FBgn0261800 | not found | 2.4286 (UP) |

[$] Song *et al.* (2).

**Table S6**: Putative Yki-Sd binding sites in regulatory regions of genes with >log$_2$2 fold change in protein levels *esg*$^{ts}$>*yki*$^{3SA}$ driven ISC tumors.

| Fly base ID | Gene Symbol | Fold enrichment in *esg*$^{ts}$>*yki*$^{3SA}$ tumors | Chromosome | $Yki-Sd binding site from TSS |
|---|---|---|---|---|
| FBgn0261276 | *opa1* | 2.0059 | 2R | -690* |
| FBgn0000562 | *egl* | 2.0314 | 2R | -9.5 |
| FBgn0035498 | *Fit1* | 2.0548 | 3L | -317 |
| FBgn0035793 | *CG7546* | 2.0985 | 3L | -533, 193 |
| FBgn0030955 | *CG6891* | 2.1608 | X | -682; -316; 456 |
| FBgn0015379 | *dod* | 3.2079 | X | -29.5 |
| FBgn0000667 | *Actn* | 2.2432 | X | 233.5 |
| FBgn0010434 | *cora* | 2.2604 | 2R | 346.5 |
| FBgn0014020 | *Rho1* | 2.2840 | 2R | 22.5; 1058.5 |
| FBgn0030052 | *CG12065* | 2.2983 | X | 87; 647.5 |
| FBgn0035499 | *Chd64* | 2.3225 | 3L | 57.5 |
| FBgn0025352 | *Mtpbeta* | 3.3272 | 2R | -129 |
| FBgn0021760 | *chb* | 2.4486 | 3L | -100; 92; 841 |
| FBgn0013733 | *shot* | 3.6429 | 2R | 94; -421.5; -1118.5; 7211; 839; 503; -114.5 |
| FBgn0033446 | *CG1648* | 2.6978 | 2R | 1867 |
| FBgn0010909 | *msn* | 3.7819 | 3L | 446; -30.5 |
| FBgn0053113 | *Rtnl1* | 3.0872 | 2L | -599.5; -308.5; -1035.5; 90.5 |
| FBgn0016726 | *Rpl-29B* | 3.1205 | 2R | -77 |
| FBgn0000043 | *Act42A* | 4.6135 | 2R | -52; -365 |
| FBgn0011225 | *jar* | 3.7014 | 3R | -562.5 |
| FBgn0028427 | *ilk* | 2.0724 | 3L | -4.5 |
| FBgn0041789 | *Pax* | 6.0551 | 2L | -19 |
| FBgn0004397 | *vinc* | 2.3977 | X | 64.5; -614 |
| FBgn0004456 | *mew* | not found | X | -185 |
| FBgn0004657 | *alphaPS2* | not found | X | 630.5; 1586; 1995.5 |
| FBgn0010395 | *Itgbeta nu* | not found | 2L | -110 |
| FBgn0002526 | *lanA* | not found | 3L | -569.5; -1244 |

$Nagaraj *et al.* (3).

**Table S7:** Change in levels of proteins in TONDU peptide-treated *esg*$^{ts}$>*yki*$^{3SA}$ driven ISC tumors.

| | | | | TREATED | |
|---|---|---|---|---|---|
| Fly base ID | Gene symbol | FDR confidence | UniProt Accession | Abundance Ratio. | T-Test (*p* value) |
| FBgn0261276 | *opa1* | High | F0JAH2 | -0.9677 | 0.0612 |
| FBgn0000562 | *egl* | Medium | Q9W1K4 | Not found | Not found |
| FBgn0260442 | *rhea* | High | Q960C2 | -0.9850 | 0.6182 |
| FBgn0035498 | *Fit1* | High | Q9VZI3 | -1.3108 | 0.4323 |
| FBgn0035793 | *CG7546* | High | M9PBU3 | -1.9655 | 0.0057 |
| FBgn0030955 | *CG6891* | High | Q8MQZ6 | -2.5940 | 0.0034 |
| FBgn0013437 | *GIP* | High | P04146 | -2.1634 | 0.0034 |
| FBgn0265991 | *Zasp52* | High | G3JX29 | -1.2146 | 0.1208 |
| FBgn0015379 | *dod* | High | P54353 | not found | Not found |
| FBgn0000667 | *Actn* | High | M9MS06 | -1.3641 | 0.0427 |
| FBgn0010434 | *cora* | High | A0A0B4LFX4 | -1.0595 | 0.0051 |
| FBgn0014020 | *Rho1* | High | P48148 | -0.8551 | 0.0536 |

| | | | | | |
|---|---|---|---|---|---|
| FBgn0030052 | *CG12065* | High | Q8MRM6 | -1.5401 | 0.0876 |
| FBgn0053470 | *CG33470; IM10; IMPPP* | High | Q8ML70 | -0.7563 | 0.4986 |
| FBgn0035499 | *Chd64* | High | M9PE30 | -1.2033 | 0.0076 |
| FBgn0025352 | *Mtpbeta* | High | O77466 | -2.3300 | 0.0139 |
| FBgn0086346 | *ALiX* | High | Q9VB05 | -2.3246 | 0.0151 |
| FBgn0260442 | *rhea* | High | M9NDM3 | -1.2592 | 0.0478 |
| FBgn0004397 | *Vinc* | High | X2JAB9 | -1.3361 | 0.0365 |
| FBgn0021760 | *chb* | High | Q9NBD7 | -2.1663 | 0.0263 |
| FBgn0262567 | *CG43107* | High | D0IQC0 | -1.6068 | 0.1094 |
| FBgn0013733 | *shot* | High | A1Z9J3 | not found | Not found |
| FBgn0033446 | *CG1648* | High | Q7K2P3 | -1.2957 | 0.0316 |
| FBgn0262735 | *Imp* | High | Q0KHU2 | -1.6114 | 0.0313 |
| FBgn0022709 | *Adk1* | High | Q9VTV3 | -1.5700 | 0.0068 |
| FBgn0086906 | *sls* | High | R4UAY6 | -1.2668 | 0.0439 |
| FBgn0010909 | *msn* | Medium | Q7KV90 | -0.9091 | 0.0273 |
| FBgn0036580 | *PDCD-5* | High | Q9VUZ8 | -1.1028 | 0.3096 |
| FBgn0051901 | *Mur29B* | Medium | Q8MS63 | -1.0776 | 0.0244 |
| FBgn0265434 | *zip* | High | A0A0B4JD95 | -1.1648 | 0.0140 |
| FBgn0005666 | *bt* | High | O76281 | -2.5538 | 0.0038 |
| FBgn0262716 | *Arp3; Arp66B* | High | P32392 | -1.1503 | 0.0234 |
| FBgn0053113 | *Rtnl1* | High | Q9VMV9 | -1.2851 | 0.0129 |
| FBgn0085282 | *CG34253* | High | A8JNV2 | -0.4406 | 0.0846 |
| FBgn0265434 | *zip-RC* | High | J7JVR0 | -1.1466 | 0.0063 |
| FBgn0016726 | *Rpl-29B* | High | B7FNL1 | -1.8229 | 0.1047 |
| FBgn0029766 | *CG15784* | High | Q9W4C1 | -2.6531 | 0.0395 |
| FBgn0038774 | *CG5023* | High | Q9I7J0 | -1.3464 | 0.0059 |
| FBgn0001217 | *Hsc70-2* | High | P11146 | -1.5196 | 0.0343 |
| FBgn0051363 | *Jupiter-RD* | High | B5RJ67 | not found | not found |
| FBgn0000043 | *Act42A* | High | P02572 | -3.6540 | 0.0297 |
| FBgn0004227 | *nonA* | High | Q8IR16 | -0.8528 | 0.0079 |
| FBgn0011225 | *jar* | High | Q01989 | -0.3926 | 0.4731 |
| FBgn0034709 | *CG3074; Swim* | High | Q7JWQ7 | -1.6677 | 0.0054 |
| FBgn0031436 | *CG3214; ND-B17.2* | High | Q9VQD7 | -0.3077 | 0.4512 |

### REFERENCES

1. Kwon Y*, et al.* (2013) The Hippo signaling pathway interactome. *Science* 342(6159):737-740.
2. Song W*, et al.* (2019) Tumor-Derived Ligands Trigger Tumor Growth and Host Wasting via Differential MEK Activation. *Dev Cell* 48(2):277-286 e276.
3. Nagaraj R*, et al.* (2012) Control of mitochondrial structure and function by the Yorkie/YAP oncogenic pathway. *Genes Dev* 26(18):2027-2037.
