## Supplementary Methods for "A *Drosophila* model of oral peptide therapeutics for adult Intestinal Stem Cell tumors"

**MATERIALS**

- Resource and Reagents

**METHODS**

- Induction of Yki-driven ISC tumors.
- Induction of somatic MARCM clones in larval wing discs.
- *UAS-vg*$^{TONDU}$ fly line.
- Design and synthesis of TONDU peptide.
- Immunostaining of the *Drosophila* adult midgut.
- EdU cell proliferation assay.
- Microscopy, image processing and GFP measure.
- Quantitative RT-PCR.
- Cell viability assay for human cancer cell lines.
- Immunofluorescence based detection of TONDU peptide in S2R+ cells.
- Immunoprecipitation studies to ascertain binding of TONDU peptide to Sd.
    - *Drosophila* cell line.
    - Plasmids used for immonoprecipitation studies
    - Protein Immunoprecipitation and Immunoblotting.
    - Chromatin Immunoprecipitation.
- Quantitation of the effect of TONDU peptide on Yki-Sd transcription using HRE-Luciferase reporter.
- Chromatin immunoprecipitation (ChIP) to determine binding of TONDU peptide to Sd in the upstream regulatory region of gene *mew*
- Proteomics of Yki-driven ISC tumors and from TONDU peptide fed flies.
    - Protein extraction from fly gut for LC-MS-MS proteomic analysis
    - Sample Preparation and Mass Spectrometric Analysis of Peptide Mixtures
    - Data Processing.
    - Proteome data analysis
    - Gene Ontology Analysis

### MATERIALS

### FLY LINES

| | Fly Genotype | Source | |
|---|---|---|---|
| 1 | *UAS-yki*$^{S111A.S168A.S250A}$ | #BDSC | #28817 |
| 2 | *UAS-yki*$^{S168A}$ | #BDSC | #28818 |
| 3 | *UAS-mew RNAi* | #BDSC | #27543 |
| 4 | *UAS-talin RNAi* | #BDSC | #28950 |
| 5 | *lgl*$^{4}$*FRT 40A* | #BDSC | #36289 |
| 6 | *esg-Gal4* | Clive Wilson, University of Oxford, UK | - |
| 7 | *UAS–torso*$^{D}$*/βcyt* | Nick Brown, University of Cambridge, UK | - |

#Bloomington Drosophila Stock Center, Indiana University, Bloomington.

### ANTIBODIES

| | Protein | Catalog Number | Source | Raised in | Working dilution |
|---|---|---|---|---|---|
| 1 | Delta (extra cellular domain) | C594.9B | #DSHB | mouse | 1:50 |
| 2 | Sd | - | Gift from (De Celis lab) | rabbit | 1:100 |
| 3 | Talin (carboxy terminus 534 amino acids) | A22A | #DSHB | mouse | 1:100 |
| 4 | αPS1 , Mew | DK.1A4 | #DSHB | mouse | 1:50 |

#Developmental Studies Hybridoma Bank, University of Iowa.

### PRIMER SEQUENCES

| | | Forward primer | Reverse primer |
|---|---|---|---|
| | **For expression analysis** | | |
| 1. | *ImpL2* | AAGAGCCGTGGACCTGGTA | TTGGTGAACTTGAGCCAGTCG |
| 2. | *yki* | CCTTGCCGCCGGGATGG | TTTGCTGCTGCTGGCGATATTG |
| 3 | *delta* | AGCGACTCTTGGTGCAGCAGGTACT | TCCGTAGTAGTTGAGATCGCAGGTGAC |
| 4 | *myc* | ACACGCGCTGCAACGATATGG | CGAGGGATTTGTGGGTAGCTTCTT |
| 5 | *wg* | TGATGGCCCTGTGCAGCG | TGATGGCCCTGTGCAGCG |
| 6 | *ex* | GCCGCCTTTACCTGTCCAAC | CGTTCCGGTTTCCAATTAGCT |
| 7 | *β-tub* | CAAGCTGGCAGTGCGGCAAC | GCTGTCACCGTGGTAGGCGCC |
| 8 | *YAP1* | ACGTTCATCTGGGACAGCAT | GTTGGGAGATGGCAAAGACA |
| | **ChIP PCR** | | |
| 9 | *Mew* | GCTTTGGTGGGGCTTGTAAC | GTAAAGGCATGAGCGCCAAAT |

**Genotype of the flies used in the study**

1. Control flies bearing GFP-marked ISCs. (Pertaining to figures 1and 4)
   *esg-Gal4, tub-Gal80$^{ts}$, UAS-GFP; +*
2. Gain of constitutively active Yki (*UAS-yki$^{S111A.S168A.S250A}$* referred to as *UAS-Yki$^{3SA}$*) in ISCs. (Pertaining to figures 1, 2 and 4; figures S1 and S3)
   *esg-Gal4, tub-Gal80$^{ts}$, UAS-GFP/+; UAS-Yki$^{3SA}$/+*
3. Simultaneous gain of constitutively active Yki (*UAS-Yki$^{3SA}$*) and TONDU peptide in ISCs. (Pertaining to figure 2; figure S2)
   *esg-Gal4, tub-Gal80$^{ts}$ ,UAS-GFP/+; UAS-Yki$^{3SA}$/+ UAS-vg$^{TONDU}$/+*
4. Downregulation of *mew* in ISCs expressing a constitutively active Yki (Pertaining to figure 4)
   *esg-Gal4, tub-Gal80$^{ts}$ ,UAS-GFP/+; UAS-Yki$^{3SA}$/+ UAS-mew RNAi/+*
5. MARCM clones with loss of *lgl* and expression of constitutively active Yki; and simultaneous gain of Yki and TONDU peptide. (Pertaining to figure S2)
   *hs-flp tub-Gal4UAS-GFP*; *lgl$^{4}$ UAS-yki$^{S168A}$ tub-Gal80 FRT40*
6. MARCM clones with loss of *lgl* and expression of constitutively active Yki and TONDU peptide.
   *hs-flp tub-Gal4UAS-GFP*; *lgl$^{4}$ UAS-yki$^{S168A}$ tub-Gal80 FRT40;UAS-vg$^{TONDU}$*
7. Gain of constitutively active βintegrin in ISCs. (Pertaining to figure S4)
   *esg-Gal4, tub-Gal80$^{ts}$, UAS-GFP/+; UAS–torso$^{D}$/βcyt*

**METHODS**

**Induction of Yki-driven ISC tumors.**

We used the UAS-Gal4 system (1) to drive constitutively active Yki (*UAS-yki$^{S111A.S168A.S250A}$*) in which 3 Serine phosphorylation sites have been mutated (2, 3), in the intestinal stem cells (ISCs), using an ISC-specific Gal4 driver (*esg-Gal4*) under control of temperature sensitive tub-Gal80$^{ts}$ (2). Flies were mated and maintained at 18$^{0}$C until eclosion of the F1 generation. Freshly eclosed F1 flies of the genotype *esg>Gal4, tub-Gal80$^{ts}$ UAS-yki$^{3SA}$* were shifted to 29$^{0}$C and maintained until dissection.

**Induction of somatic MARCM clones in larval wing discs.**

Somatic clones with loss of *lgl* and gain of Yki; or loss of *lgl* with simultaneous gain of Yki and TONDU were generated by MARCM technique (4). Briefly, heat shock was given to synchronized early second instar larvae at 48 hours after egg laying at 37°C for 15 min. Larvae were maintained at $25^0$C and dissected 96 hours after heat shock.

**Generation of *UAS-vg$^{TONDU}$* fly line**

We synthesized oligonucleotide coding for the *Drosophila* TONDU domain (CVVFTNYSGDTASQVDEHFSRALNY) (5). We introduced a start (ATG) and stop codon (TAA) flanking the nucleotide sequences, and inserted a 5'EcoR1 and 3' Xba1 endonuclease restriction enzyme site on either side to allow directional cloning into pUASt vector (Addgene). We replaced Cytosine on position one and Alanine on position 22 with Serine (SVVFTNYSGDTASQVDEHFSRSLNY) to make the encoded peptide more polar and therefore improve its solubility. Substitution of terminal Cysteine would also reduce chances of aberrant dimer formation. The VXXHF domain of the TONDU domain, which is essential for interaction with TEAD/Sd (6), was left unchanged. The synthesized oligo was cloned into pUAST vector carrying *mini white*, and injected into embryos of CS flies at C-CAMP (Center for Cellular and Molecular Platforms, NCBS, Bangalore, India). Adults were screened for insertion of the vector into the third chromosome.

**Design and synthesis of the TONDU peptide and its variants.**

**TONDU-peptide**: We synthesized a peptide corresponding to the TONDU domain with certain modifications. The basic peptide is a 46 amino acid peptide (YGRKKRRQRRRGGPKKKRKVGG [VVFTNYSGDTASQVDEHFSRALNY]) comprised of a 24 aa of the TONDU domain (VVFTNYSGDTASQVDEHFSRALNY) preceded by the conserved SV40 T-Antigen nuclear localizing signal (PKKKRKV) (7) and a cell penetrating peptide (YGRKKRRQRRR) derived from human immunodeficiency virus (HIV) (8), and a NLS sequence flanked by a di-glycine (GG) spacer to avoid any steric hindrance between a tag and the rest of the peptide. The first Cytosine on the TONDU domain was removed to prevent dimerization of the peptide.

**FLAG-tagged TONDU peptide**: To test for binding partners to TONDU peptide, we added a FLAG tag (DYKDDDDK) at its C-terminus (YGRKKRRQRRRGGPKKKRKVGG-

VVFTNYSGDTASQVDEHFSRALNYDYKDDDDK) to allow protein immunoprecipitation using an anti-FLAG antibody.

**Fluorescent-tagged TONDU peptide:** To track uptake of the peptide and facilitate its cellular localization, we added a 5-TAMARA, a fluorescent tag, to the C-terminus of the TAT-NLS-TONDU peptide. The peptides were synthesized at GL Biochem Shanghai Ltd.

**Immunostaining of the *Drosophila* adult midgut**

Prior to dissection, female flies of desired genotype were starved briefly and fed water for 2 hours to flush out food from the gut. Midguts were dissected in 1X PBS and fixed in 4% paraformaldehyde in PBS containing 0.2% triton X-100 for 30 minutes at room temperature; followed by washing in PBS containing 0.2% triton X-100 for 15 minutes. The guts were then incubated in primary antibody (see table below) at $4^0$C overnight, followed by blocking with 0.1 % BSA for 1 hour and incubation with secondary antibody (Alexa fluor 555) for 4 hours at room temperature. Next, the guts were washed in 1X PBS followed by mounting in an anti-fade mounting medium, Vectashield (Sigma), and counterstained for nuclei using TOPRO (Invitrogen, S33025) or F-actin using Alexa Fluor Phalloidin-633 (Invitrogen A22284, 1:100).

**Microscopy and Image Processing**

Images were acquired using a Leica SP5 confocal microscope and processed using the Leica application software and Adobe Photoshop CS5.

**Measurement of GFP from confocal images.**

GFP was quantitated from full projections of images acquired using confocal microscopy. GFP intensity in grayscale from ROIs covering the entire gut was acquired using the Leica-LSM proprietary software. The GFP intensity was normalized to the area of each ROI. Student t-test was done using MS-Excel to look for statistical significance in GFP variation. Box plots were generated using GraphPad Prism 7.0.

**EdU cell proliferation assay**

Cell proliferation was detected by EdU uptake using Click-iT Alexa-Fluor-555 290 kit by Invitrogen. Briefly, unfixed guts from female $esg^{ts}$>$UAS\text{-}yki^{3SA}$flies were incubated with100 μM of EdU in Schneider's insect medium, for one hour at room temperature. Tissue was then fixed in 4% paraformaldehyde and incubated in secondary buffer containing fluorescent-tagged dye (following manufacturer's instruction) for one hour at room temperature and subsequently washed in PBS, counter-stained with TO-PRO-3 (Invitrogen, S33025) and mounted using an anti-fade mounting medium (Invitrogen).

**Quantitative RT-PCR**

RT-PCR was performed using SYBR green (Applied Biosystems) on ABI7 900 HT. Prior to dissection, female $esg^{ts}$>$UAS$-$yki^{3SA}$ flies were starved briefly and fed water for 2 hours to flush out food from the gut. Total RNA from 20 midguts was isolated using Qiagen RNeasy columns. RNA was treated with RNase-free DNase (Roche), to get rid of any traces of DNA, before converting RNA to cDNA using a cDNA preparation kit (Invitrogen). The resulting cDNA was used as substrate for relative quantitation using SYBR green on ABI7 900 HT. β-Tubulin was used as an endogenous control. Genes were assayed from four biological replicates for each condition. qPCR was performed using the following conditions: DNA polymerase activation for 10 min at 95°C, followed by 40 cycles of duplex melting for 15 s at 95°C and a combined annealing and extension step for 1 min at 60°C. The threshold-cycle (Ct) values were generated automatically. The relative expression value of each gene in the two conditions was calculated using the 2-ΔΔCt method.

**Cancer cell line and cell culture conditions**

The prostate (PC3 and LNCaP) and colorectal (Colo 320-HSR) cancer cell lines were obtained from American Type Cell Culture (ATCC, Manassas, VA, USA). The colorectal cancer cell line WiDr was a kind gift from Dr. Eric R. Fearon, University of Michigan, Ann Arbor, MI, USA. All of the cell lines were cultured as per ATCC guidelines in a $CO_2$ incubator (Thermo-Fisher) supplied with 5% $CO_2$ at 37°C temperature. Cell line authentication was done via short tandem repeats (STR) profiling at Lifecode Technologies Private Limited (Bangalore, India) and DNA Forensics Laboratory (New Delhi, India). Routine check for mycoplasma contamination of all cell lines was carried out using PlasmoTest mycoplasma detection kit (InvivoGen).

**Cell viability assay of human cancer cell lines**

To determine the effect of TONDU peptides on the cell viability of prostate cancer (PC3 and LNCaP) and colorectal cancer (COLO320 and WiDR) cells. Approximately 3,000 cells were plated in each well of a 96-well plate. After 24 hours, TONDU peptide was added to the cultured cells at three different concentrations: 50 nM, 100 nM and 250 nM. No peptide was added in the control group. After 72 and 96 hours of peptide treatment, cell viability was determined using resazurin sodium salt solution (R7107, Sigma). Briefly, resazurin (0.02mg/ml; w/v) diluted in culture media was added to the cells and incubated for 4 hours in dark at 37ºC. The fluorescence was measured at 530/590 nm (excitation/emission) using BioTek™ Synergy™ H4 Hybrid Microplate Reader (Winooski, VT, USA).

*Statistical significance*

Biologically independent samples were used (n=3) in each experiment; data represents mean ± SEM. Statistical significance was determined using two-tailed unpaired Student's t-test, $*P \leq 0.05$ and $**P \leq$ *0.001.*

**Immunoprecipitation studies to ascertain binding of TONDU peptide to Sd**

- ***Drosophila* cell line**

  *Drosophila* S2R+ cells (sex: male) were cultured in Schneider's medium supplemented with 10% fetal bovine serum (FBS) at 25°C.

- **Plasmids used for immonoprecipitation studies**

  The full-length cDNAs of Sd (GEO03367) and Yki (GEO02945) from the Drosophila Genomics Resource Center were cloned into the Drosophila Gateway vector pAWH and pAWG respectively. GFP was cloned into pAWM as a control.

- **Immunoprecipitation and Immunoblotting** were performed as previously described (9). In brief, DNA was transfected into S2R+ using Effectene transfection reagent (Qiagen, 301427). After 2 days of incubation, cells were incubated with or without 1 µM TONDU peptide for 24 hrs and then lysed with lysis buffer (Pierce 87788) containing a protease and phosphatase inhibitor cocktail (Pierce, 78440). Lysate was incubated with Chromotek-GFP-Trap (Bulldog Biotechnology, gta-20) for 2 hr at 4°C to precipitate the proteins. Beads were washed 3-4 times with 1 mL lysis buffer and then then boiled in SDS sample buffer, run on a 4%–20% polyacrylamide gel (Bio-Rad, 4561096), and transferred to an Immobilon-P polyvinylidene fluoride (PVDF) membrane (Millipore).

The membrane was blocked by 5% BSA in TBST (TBS with 0.1% Tween-20) in room temperature for 1 hr and then probed with anti-GFP (Molecular Probes, A6455), anti-HA (Covance/BioLegend, MMS-101P), or anti-FLAG (Sigma, F3165) antibody in 1X TBST with 5% BSA overnight, followed by HRP-conjugated secondary antibody, and signal was detected by enhanced chemiluminescence (ECL; Amersham, RPN2209; Pierce, 34095).

- For the TONDU-Sd binding assay, HA-Sd was expressed in S2R+ cells and purified through immunoprecipitation with RIPA buffer (Pierce, 89901) and anti-HA agarose (Sigma, A2095). Purified HA-Sd proteins were incubated with 1 µM TONDU peptide directly. The sample was then washed and subjected to immunoblotting.

**Quantitation of effect of TONDU peptide on Yki-Sd-driven transcription using HRE-Luciferase reporter**

*Drosophila* S2R+ cells were maintained at 25$^0$C in Schneider's medium (GIBCO) with 10% heat-inactivated fetal bovine serum (Sigma) and 5% Pen-Strep (GIBCO). Experiments were run in 24-well plate, three replicates per condition. Cells were co-transfected with 100 ng each of 1) HRE-luciferase reporter (containing two copies of a Hippo Response Element cloned upstream of a hsp70 basal promoter in pGL3 basic vector (10)), along with 2) Sd- or 3) Yki-expressing pAc5.1/V5-HisB plasmids (10) (gift from D. Pan); 10 ng of Act-Renilla was used for transfection control. Transfection was carried out using Effectene (Qiagen), as per manufacturer's recommended protocol. 24 hours after transfection, 50 or 100 nM of the TONDU peptide was added to wells in triplicate. 48 hours after addition of the TONDU peptide, cells were harvested and Luciferase activity was measured using Dual Glo (Promega) as per the kit instructions, and measured using a Spectramax Luminescence plate reader.

**Detection of fluorescent labeled TONDU peptide in S2R+ cells**

*Drosophila* S2R+ cells were grown to confluence in Schneider's medium (GIBCO) supplemented with 10% heat-inactivated fetal bovine serum (Sigma), and 5% Pen-Strep (GIBCO) at 25$^0$C in 24 well plates. TAMARA-tagged TONDU peptide was added to the medium to a final concentration of 100 nM and cells were incubated for 6 hours. Next, the medium was discarded and cells washed 3 times with 1XPBS. The cells were then taken and put on lysine coated slides, fixed with 4% formaldehyde in 1XPBS, and counterstained with DAPI. Cells were imaged with a Nikon Ti, CSU-X1 spinning disk confocal microscope and the images were processed using Fiji image processing software (https://imagej.net › Fiji).

**Chromatin immunoprecipitation (ChIP) to determine binding of TONDU peptide to Sd in the upstream regulatory region of gene *mew***

ChIP was performed using LowCell# ChIP kit protein A (Diagenode Cat# C01010072) according to manufacturer's instructions. Briefly, midguts from 35 adult female *esg*$^{ts}$>*UAS-yki*$^{3SA}$ flies (pre-starved for 1 hour) were dissected in ice-cold 1XPBS and crosslinked in 1% formaldehyde (Sigma) for 15 minutes at 37$^0$C. Crosslinking was quenched with 125 mM Glycine. The guts were washed with PBS and precipitated with centrifugation at 6000 rpm for 5 minutes. The pellet was lysed in 250 mL of Buffer B (LowCell# ChIP kit) supplemented with complete protease inhibitor (Roche) and PMSF (Sigma). 130 µL of lysed chromatin was sheared using a Bioruptor (Diagenode) at high frequency for 15 cycles of 30 sec ON, 30 sec OFF. 870 µL of Buffer A (LowCell# ChIP kit) supplemented with complete protease inhibitor (Roche) and PMSF (Sigma) was added to the shared chromatin. 8 µL of the chromatin solution was saved as an input control. 11 µL of magnetic beads was washed twice with Buffer-A (LowCell# ChIP kit) and resuspended in 800 µL of Buffer A. 2 µg of Anti-FLAG antibody (Sigma, F1804) were added to the washed beads and gently agitated at 4$^0$C for 4 hours. The beads-antibody complex was precipitated with a magnet and the supernatant was removed. 800 µL of shared chromatin was added to the beads-antibody complex and rotated at 4$^0$C overnight. The immobilized chromatin was then washed with Buffer A three times and Buffer C once, and eluted in 100 µL elution buffer (1% SDS, 0.1 M sodium bicarbonate with proteinase K and RNaseA). The chromatin was subjected to either phenol-chloroform extraction for DNA purification and subsequent qPCR analysis; or the protein or the protein was extracted by heating the washed beads at 95$^0$C in 20 ul SDS loading dye (4X) for 10 minutes and centrifuged at 13000 rpm for 10 minutes. The supernatant was collected and used for dot blot analysis.

**Protein dot blot**

1 mM TONDU peptide was serially diluted ($10^{-1}$, $10^{-2}$, $10^{-3}$) and blotted using a narrow-mouth pipette tip, and 7.5 µl of peptide or enriched protein fraction from ChIP were applied slowly onto the nitrocellulose membrane (Thermo Fisher, 0.2 um pore size). The membrane was air dried and

then blocked in 5% BSA in TBS-T (Tris-buffered saline, 0.1% Tween 20) for 2 hours at RT, then incubated for 3 hours with a secondary antibody conjugated with HRP (Jackson Immuno Research #711035152), washed three times with TBST, and then detected with chemiluminescent substrate (Thermofischer #34080) and visualized on X-ray film (Fuji, Super HR-t).

**Proteomics of Yki-driven ISC tumors**

- **Protein extraction from fly guts for LC-MS-MS analysis**

Prior to dissection, female $esg^{ts}$>*UAS-yki*$^{3S}$ flies were briefly starved and fed on water for 2 hours to clear the gut. Adult guts were dissected in cold 1X PBS from 20 flies. The fore- and hindguts were removed, and the midguts were put in 100 µl extraction buffer (6M GnHCl in 50mM Tris-Cl pH 7.4, 65 mM DTT) with 50 mM sodium acetate and protease inhibitors (1X protease inhibitor cocktail with 0.2 mM PMSF) was added to the sample. The guts were sonicated with a Bioruptor (Diagenode) using the following settings: Sonication cycle: 30 sec ON and 30 sec OFF for 5 cycles at 4˚C. The cell debris was removed by brief centrifuging it at 8000 RPM for 3 minutes; then supernatant was transferred to new tube. The protein concentration was determined spectrophotometrically using Nanodrop and using BCA protein assay (Thermo fisher) following the manufacturer’s protocol. 5 µg of the protein was used for LC-MS-MS analysis. We made certain that the tissue was processed within half an hour of dissection.

**Sample preparation for LC-MS/MS**

5 µg of the protein samples were reduced with 5 mM TCEP, further alkylated with 50 mM iodoacetamide, and digested with Trypsin (1:50, Trypsin/lysate ratio) for 16 hours at 37°C. Digests were cleaned using a C18 silica cartridge to remove the salt and dried using a speed vac. The dried pellet was resuspended in 5% acetonitrile, 0.1% formic acid (Buffer A).

**Mass Spectrometric Analysis of Peptide Mixtures**

The experiment was performed using an EASY-nLC 1000 system (Thermo Fisher Scientific) coupled to a Thermo Fisher-Orbitrap *Fusion* mass spectrometer equipped with a nanoelectrospray ion source. 1.0 µg of the peptide mixture was resolved using a 25 cm Thermo Easy-spray PepMap C18 column. The peptides were loaded with Buffer A and eluted with a 0–

40% gradient of Buffer B (95% acetonitrile, 0.1% formic acid) at a flow rate of 300 nl/min for 60 min. MS data was acquired using a data-dependent top20 method dynamically choosing the most abundant precursor ions from the survey scan.

**Data Processing**

All samples were processed and the 8 RAW files generated were analyzed with Proteome Discoverer (v2.2) against the Uniprot *Drosophila melanogaster* reference proteome database. For Sequest search, the precursor and fragment mass tolerances were set at 10 ppm and 0.5 Da, respectively. The protease used to generate peptides, *i.e.* enzyme specificity, was set for trypsin/P (cleavage at the C terminus of "K/R: unless followed by "P") along with a maximum missed cleavages value of two. Carbamidomethyl on cysteine as fixed modification and oxidation of methionine and N-terminal acetylation were considered as variable modifications for database search. Both the peptide spectrum match and the protein false discovery rate were set to 0.01 FDR.

**Proteome data analysis**

To look for biologically relevant protein signatures in $esg^{ts}$>$yki^{3SA}$ tumors and also to look for their status in the presence of the TONDU peptide, we calculated the log2 abundance ratios, using mean abundance values for individual Uniprot IDs of $esg^{ts}$>*UAS*-$yki^{3SA}$ day 7 versus day 1 proteome. Only those were taken into consideration whose combined FDR confidence was <0.05 (medium) or <0.01 (high), while those with >0.05 (low) were discarded. We further filtered out those peptides that were not detected in either MS, or MS-MS spectrum, depending on its peak calling. We noted the number of peptides that matched each Uniprot ID, ranged from 1 to 67. To ascertain statistically significant calls, we applied student t-test on replicate readings for the individual Uniprot IDs and only those with $P<0.05$ were considered. We first calculated log2 abundance ratio of proteins in day 7 with day 1 of $esg^{ts}$>$yki^{3SA}$ tumors, and considered only those gene products whose log2 fold change was ≥ 2.  Similarly we calculated the log2 abundance ratio of proteins from peptide treated and untreated $esg^{ts}$>$yki^{3SA}$ tumors. For which we included two replicates of $esg^{ts}$>$yki^{3SA}$ flies fed on 200 µM of the TONDU peptide and one replicate from $esg^{ts}$>$yki^{3SA}$ *UAS*-$vg^{TONDU}$; which were compared to day 7 $esg^{ts}$>$yki^{3SA}$ tumors from unfed flies.

We applied the Student t-test to look for statistical significance for each log2 fold change, and considered only those with $P<0.05$.

### Gene Ontology Analysis

To identify biological function of genes and look for enrichment of functional classes, we undertook Gene Ontology analysis using PANTHER (Protein ANalysis THrough Evolutionary Relationships) classification system (http://www.pantherdb.org, (11). Protein functions were inferred by classification of genes into one or more groups, depending on 1) Molecular function 2) Biological Process m 3) Protein class 4) Pathways and 5) Cellular component.

**Heat Maps:** Heat maps were generated using Heatmapper (http://heatmapper.ca/). For the heatmap in Fig 3A, raw abundance values for individual UniProt IDs were subjected to row scaling, and clustered using average linkage clustering with Euclidean method for distance measure.

### REFERENCES


1. Brand AH & Perrimon N (1993) Targeted gene expression as a means of altering cell fates and generating dominant phenotypes. *Development* 118(2):401-415.
2. Kwon Y*, et al.* (2015) Systemic organ wasting induced by localized expression of the secreted insulin/IGF antagonist ImpL2. *Dev Cell* 33(1):36-46.
3. Oh H & Irvine KD (2009) In vivo analysis of Yorkie phosphorylation sites. *Oncogene* 28(17):1916-1927.
4. Lee T & Luo L (2001) Mosaic analysis with a repressible cell marker (MARCM) for Drosophila neural development. *Trends Neurosci* 24(5):251-254.
5. Pobbati AV & Hong W (2013) Emerging roles of TEAD transcription factors and its coactivators in cancers. *Cancer Biol Ther* 14(5):390-398.
6. Pobbati AV, Chan SW, Lee I, Song H, & Hong W (2012) Structural and functional similarity between the Vgll1-TEAD and the YAP-TEAD complexes. *Structure* 20(7):1135-1140.
7. Lanford RE, Kanda P, & Kennedy RC (1986) Induction of nuclear transport with a synthetic peptide homologous to the SV40 T antigen transport signal. *Cell* 46(4):575-582.
8. Vives E, Brodin P, & Lebleu B (1997) A truncated HIV-1 Tat protein basic domain rapidly translocates through the plasma membrane and accumulates in the cell nucleus. *J Biol Chem* 272(25):16010-16017.
9. Tang HW*, et al.* (2018) The TORC1-Regulated CPA Complex Rewires an RNA Processing Network to Drive Autophagy and Metabolic Reprogramming. *Cell Metab* 27(5):1040-1054 e1048.
10. Wu S, Liu Y, Zheng Y, Dong J, & Pan D (2008) The TEAD/TEF family protein Scalloped mediates transcriptional output of the Hippo growth-regulatory pathway. *Dev Cell* 14(3):388-398.

11. Mi H, Guo N, Kejariwal A, & Thomas PD (2007) PANTHER version 6: protein sequence and function evolution data with expanded representation of biological pathways. *Nucleic Acids Res* 35(Database issue):D247-252.
